# Endothelial expression of ZBTB16 protects against cardiac aging

**DOI:** 10.1101/2025.10.08.681100

**Authors:** Kathrin A. Stilz, Vincent Elvin Leonard, David Rodriguez Morales, Simone-Franziska Glaser, Veronica Larcher, Mariano Ruz Jurado, Pedro Felipe Malacarne, Nivethitha Manickam, Lukas S. Tombor, Wesley Abplanalp, Josefine Panthel, Haris Kujundzic, Ariane Fischer, Katja Schmitz, Oliver J. Müller, Susanne Hille, Christian Kupatt, Tarik Bozoglu, Haider Sami, Manfred Ogris, Tara Procida-Kowalski, Marek Bartkuhn, David John, Michail Yekelchyk, Tessa Schmachtel, Michael Rieger, Minh-Duc Pham, Jaya Krishnan, Stefan Günther, Ralf P. Brandes, Thomas Braun, Andreas M. Zeiher, Julian U. G. Wagner, Stefanie Dimmeler

## Abstract

**Background and aim:** Aging significantly increases the risk of cardiovascular diseases, characterized by progressive cardiac dysfunction. The vascular niche is crucial for maintaining cardiac homeostasis, yet endothelial cell (EC) impairment during aging remains poorly understood. This study investigates epigenetically regulated mechanisms mediating EC-dependent cardiac aging and identifies a critical role of Zinc finger and BTB domain-containing protein 16 (ZBTB16).

**Methods:** Chromatin accessibility (snATAC-seq) and transcriptomic (snRNA-seq) analyses were performed on aged hearts to identify age-related regulatory changes. Functional studies using genetic models, assessed cardiac aging phenotypes. In vitro assays examined EC senescence and secretory profiles, while co-culture experiments analyzed the impact of ZBTB16-deficient EC supernatants on fibroblasts, cardiomyocytes, and neurons. Overexpression experiments in vitro and in vivo tested the potential for ZBTB16 to mitigate aging-associated dysfunction.

**Results:** Aged hearts exhibited decreased chromatin accessibility and expression of the transcription factor ZBTB16 in both human and mice. Loss of ZBTB16 in young mice, including *Zbtb16* haploinsufficient and endothelial-specific knockout mice, led to premature aging, diastolic dysfunction, and increased secretion of pro-fibrotic and inflammatory factors. Supernatants from ZBTB16-deficient ECs activated fibroblasts, induced cardiomyocyte hypertrophy, and impaired neuronal sprouting. Overexpression of ZBTB16 reversed these effects in senescent ECs and aged mice and reduced diastolic dysfunction. Mechanistic studies identified key downstream targets of ZBTB16, including nuclear receptor-interacting protein 1 (NRIP1). ZBTB16 suppressed NRIP1 expression, limiting fibroblast activation and pro-fibrotic signaling.

**Conclusions:** ZBTB16 is a key regulator of endothelial function, maintaining vascular niche homeostasis and mitigating aging-associated cardiac dysfunction. Its loss promotes EC senescence and pro-fibrotic signaling, contributing to diastolic dysfunction. Overexpression of ZBTB16 presents a potential therapeutic strategy for preserving cardiac function during aging. These findings establish a novel role for ZBTB16 in endothelial aging and cardiovascular disease prevention.

**Graphical Abstract:** 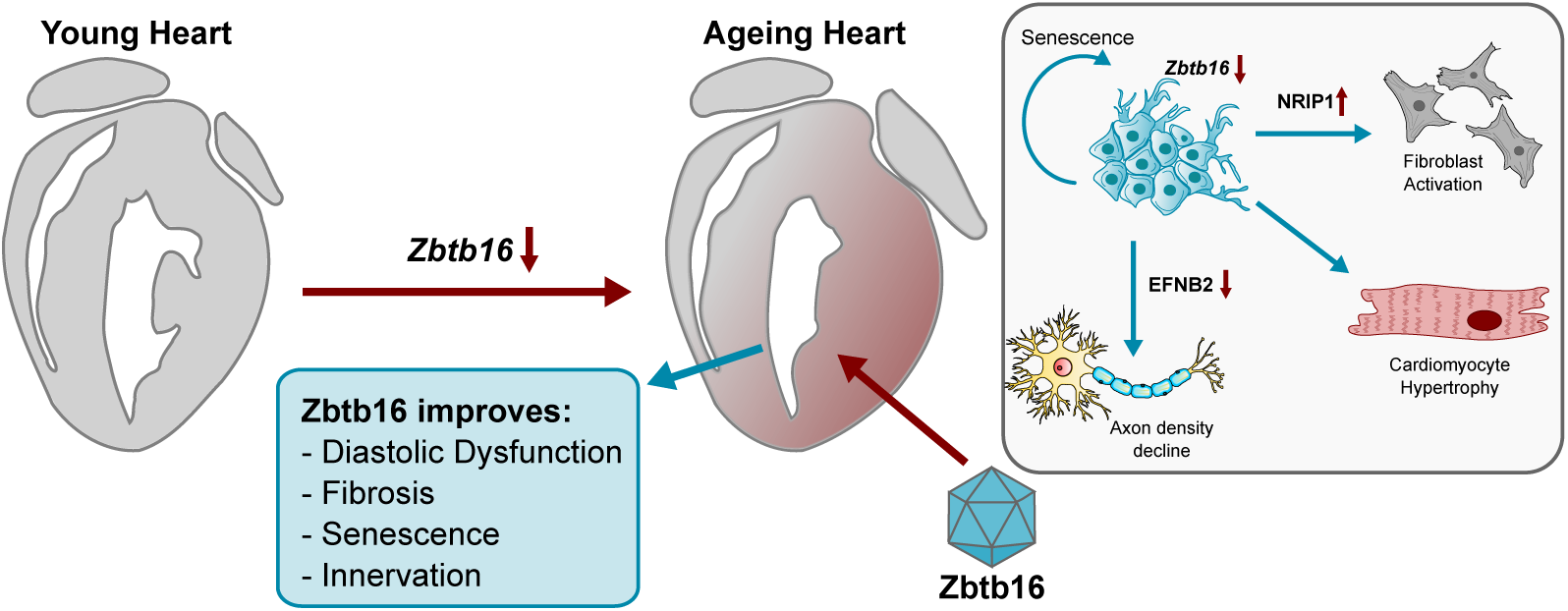

## INTRODUCTION

Life expectancy has significantly increased over the past century. A longer lifespan, however, comes with an increased risk for age-associated diseases, such as cardiovascular disease, which is still the leading cause of death worldwide^1^. Increasing age reduces diastolic heart function accompanied by increased cardiac fibrosis and microcirculatory dysfunction^2^. Reduced microcirculatory function can limit oxygen and nutrient supply. Aging and induction of endothelial cell senescence additionally affects the secretion of endothelial-derived paracrine factors, which are important to control organ homeostasis, repair and regeneration. Accumulation of senescent endothelial cells and reduced vasodilation have been associated with heart failure with preserved ejection fraction, which is the typically age-associated manifestation of heart failure^3^. Recent studies suggest that improving EC functions can prevent many of the age-associated impairments of organ function^4^. Likewise, senolytic treatments, which remove senescent cells, show promising benefits in cardiovascular disease models^5–10^.

On a molecular level, endothelial cell senescence is well known to reduce EC nitric oxide synthesis associated with an altered redox balance^7^. Aging further induces proinflammatory senescence associated secretory proteins (SASPs), activates the expression of endothelial adhesion molecules (e.g. ICAM-1), augments prothrombotic metalloprotease activity, while increasing the production of Angiotensin II, endothelin 1 and IGF binding protein expression^7^. However, general up-stream epigenetic and transcriptional regulators, which govern the age-associated changes in endothelial cells are less known. It has been shown that activation of NFkB contributes to endothelial inflammation during aging^11^, but a systematic analysis of epigenetic regulation of endothelial cells during aging in vivo is missing.

To gain insights into how gene activity and accessibility are regulated in aging cardiac endothelial cells, we investigated chromatin accessibility, which is controlled by epigenetic changes such as histone modifications or DNA methylation, and the transcriptional regulation, which together controls gene expression. For this, we employed single-nucleus ATAC sequencing in young and old mice. Our study revealed that the Krueppel C2H2-type zinc-finger protein ZBTB16 was among the top repressed genes. ZBTB16 originally captured attention as translocations with the retinoic acid receptor locus generated a fusion protein, which contributes to the development of leukemia^12^. Apart from its involvement in leukemia, ZBTB16 has been implicated in various cellular processes, including the regulation of cell growth, apoptosis (programmed cell death), and differentiation^13–15^. ZBTB16 also was shown to modulate inflammatory and antiviral immune responses as a transcription factor by directly inducing gene expression via its C-terminal Krueppel-type zinc-fingers^16^. The N-terminal BTB/POZ and RD2 domains can additionally recruiting other factors thereby reducing TLR-dependent activation of inflammatory genes^17^. However, recently also pro-inflammatory properties have been described: ZBTB16 was shown to activate the assembly of the inflammasome by inducing sumoylation of the inflammasome component ASC (apoptosis-associated speck-like protein containing a CARD)^18^. While the specific functions of ZBTB16 in the endothelium in vivo has not been explored, in vitro studies suggest a pro-proliferative^19^ and autophagy-inducing function^20^.

Here we investigated the function of the epigenetically repressed ZBTB16 in cardiac aging in mice.

## MATERIAL AND METHODS

Descriptions of the individual methods and the source of the respective reagents are described in the *Extended Material and Methods* section. The key animal experiments are described below:

### Laboratory animals and in vivo experiments

C57Bl/6J wildtype mice were purchased from Janvier (Le Genest SaintIsle, France) and from Charles River (Sulzfeld, Germany). Homozygosity of these inbred mice was controlled by Janvier and Charles River using exome sequencing. Zbtb16-deletion mice (Strain ID: 066990-UCD) were purchased from UC Davis MMRRC (Los Angelos, CA, USA). Zbtb16fl/fl mice were purchased from GemPharma Tech (strain number: T016043). All animal experiments have been executed in accordance to the guidelines from Directive 2010/63/EU of the European Parliament on the protection of animals used for scientific purposes and were approved by the local authorities by the state of Hesse (Regierungspräsidium Darmstadt). Experiments were conducted as follows: mice were anaesthetized with 2-2.5% of isoflurane and echocardiography was performed to monitor heart function using the Vevo 3100 echocardiography system with the Vevo LAB software (Fujifilm VisualSonics). Systolic function (ejection fraction) and diastolic function (expressed as E/E’ * (-1)) were measured. At the end of the experiment, mice were anaesthetized with 2-2.5% of isoflurane and euthanized via cervical dislocation.

### AAV9-mediated Zbtb16 induction

To induce endothelial-targeted *Zbtb16* expression, AAV9 vectors encoding the murine *Zbtb16* or fLuc2 under the control of the *endoglin* promotor were generated. Viral particles were coated with a G2-PAMAM linked to an endothelial-targeted peptide (PAMAM-G2^CNN^) which were generated as previously described^21,22^ and referred to as EC-AAV9-Zbtb16. Viral particles were diluted in OptiMEM GlutaMAX^TM^ (51985026, Gibco) and coated with PAMAM G2CNN (1.8 µg of PAMAM-G2^CNN^ to coat 2×10^12^ viral particles) for 30 minutes at room temperature, prior to injection. 2×10^12^ AAV9-Zbtb16 were i.v. injected to 18 months old C57Bl/6J mice. AAV9 encoding for fLuc2 served as negative control.

### Statistical analysis

Statistical data are represented as mean and error bars indicate the standard error of mean (SEM). Shapiro Wilk test was carried out the normality distribution analysis. To compare two groups, an unpaired two-sided t-test (Gaussian distributed data) or a Mann Whitney U test (non-Gaussian distributed data) was used. For multi-group comparison, an ordinary one-way ANOVA with a post-hoc Tukey or Dunnett’s test (Gaussian distributed data) or a Kruskal-Wallis test (non-Gaussian distributed data) was used.

## RESULTS

### Epigenetic alterations in aged cardiac endothelial cells

To investigate the epigenetic control of cardiac endothelial cell aging, we analyzed chromatin accessibility using single nuclei assay for transposase-accessible chromatin (snATAC-Seq) in hearts from young (3 months) and old (22 months) mice. Unsupervised clustering of the snATAC-seq data revealed six distinct cellular populations: cardiomyocytes, endothelial cells, epicardial cells, fibroblasts, pericytes, and purkinje cells (**Fig. 1A, Supplementary data online Fig. 1**). A detailed analysis of the endothelial cluster identified 383 differentially accessible regions, of which 197 exhibited reduced and 186 showed increased accessibility in aged endothelial cells (**Fig. 1B**). Genes with increased accessibility were linked to processes such as “cell-cell adhesion,” “synaptic signaling,” and “neuroprojection” (**Supplementary data online Fig. 2A**). Notable genes included *Robo2, Slit3*, and *Cadps2*, which control synaptic development and cardiac innervation (**Supplementary data online Fig. 2B**). Additional genes like *Sgpl1* (implicated in endothelial barrier dysfunction) and solute carriers *Slc6a6* and *Slc7a11* were also upregulated (**Supplementary data online Fig. 2C)**. Conversely, reduced chromatin accessibility was associated with genes involved in “myofibril assembly” and “regulation of actin-filament-based processes,” such as *Actn2*, *Troponin*, and *Fhl2* (**Supplementary data online Fig. 2D-E**).

**Figure 1:**
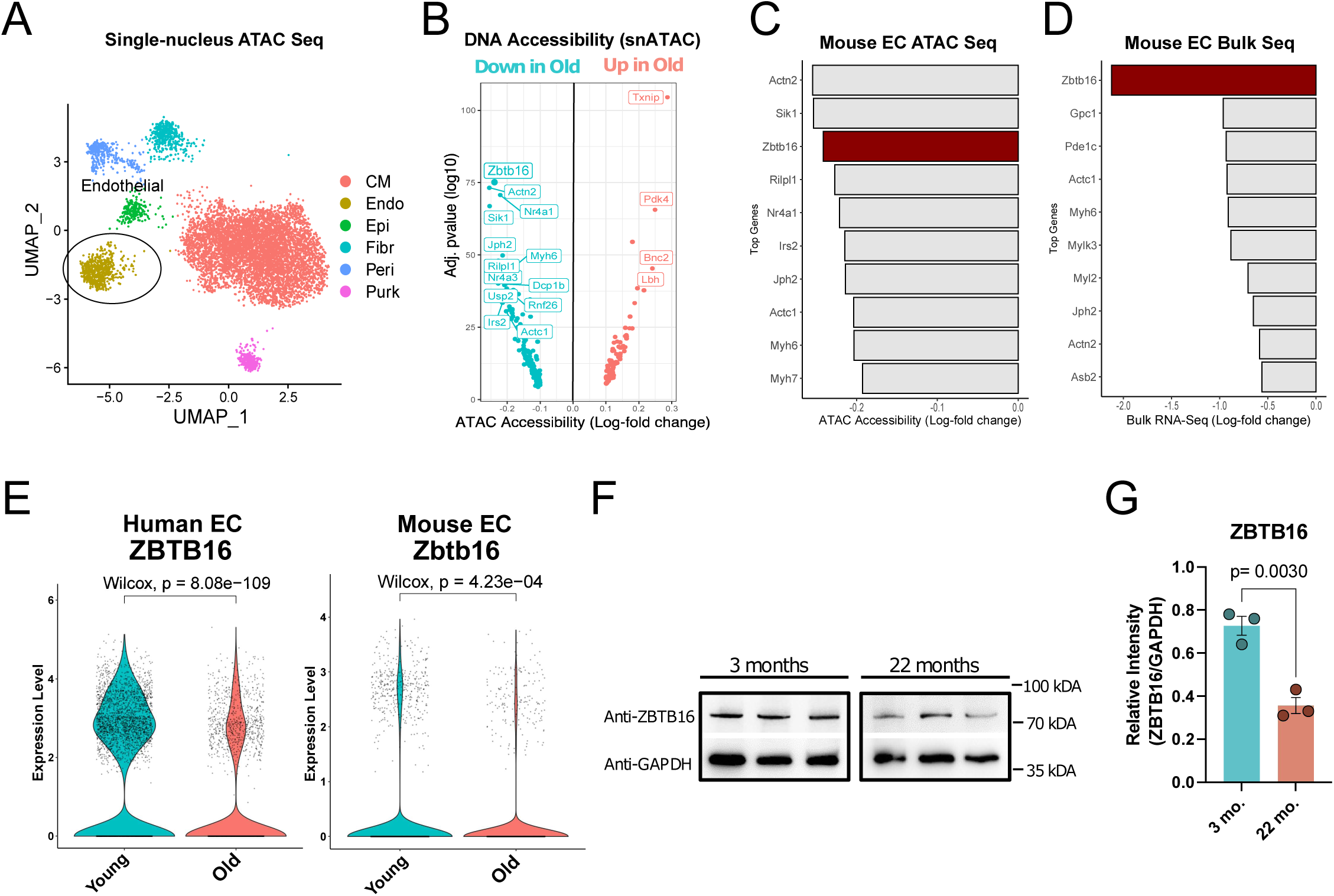
*Zbtb16* is repressed in aged cardiac endothelial cells. (**A**) UMAP representation of single-nucleus ATAC sequencing data from young (3 months) and aged (22 months) mice (n=2). Colors depict major cardiovascular cell types. (**B**) Volcano plot showing top differential (Log-fold change > 0.1 or < -0.1) significant (Bonferroni adj. p-value <0.05) accessible genes in young and old endothelial cells in single nucleus ATAC sequencing. (**C**) Bar plot showing top 10 less accessible genes (ATAC) in aged ECs among the intersection of significant regulated genes in mouse EC bulk RNA sequencing of young and old mice (n=3). (**D**) Bar plot of top 10 downregulated genes in mouse EC bulk RNA sequencing of young (3 months) and aged (20 months) mice (n=6). (**E**) right: Violin Plot of ZBTB16 expression in human ECs. Data show individuals < 55 years (blue, young) and > 55 years (red, old). Data taken from heartcellatlas.org. Bonferroni adj. p-value, Wilcox test. Left: Violin Plot of ZBTB16 expression in young and old mouse ECs. Data taken from^8^. (**F, G**) Western Blot showing ZBTB16 expression in young (3 months) and old (22 months) whole mouse heart cell lysates (n=3). Quantification is shown in G. Data was normalized to GAPDH loading control band intensity. Data are represented as mean and error bar indicate the standard error of the mean. P-value was calculated by two-tailed Student’s t-test.

Among the downregulated genes, the zinc finger transcription factor *Zbtb16* stood out. It was the only gene that overlapped among the top differentially accessible and differentially expressed genes in bulk RNA sequencing data from isolated endothelial cells from young and aged mice (**Fig. 1C-D, Supplementary data online Fig. 3**). Validation using single-nuclei RNA sequencing confirmed *ZBTB16* downregulation in aged human and mouse cardiac tissues, respectively (**Fig. 1E**). Additional experiments, including in situ hybridization and immunoblotting, corroborated these findings, demonstrating reduced *Zbtb16* expression at mRNA and protein levels in aged mouse hearts (**Fig. 1F, G, Supplementary data online Fig. 4**). In addition, we found a significant down-regulation of ZBTB16 mRNA expression in endothelial cells and other cell types in hearts from patients with heart failure with preserved ejection fraction and cardiac hypertrophy (**Supplementary data online Figure 5**, data taken from^23,24^).

### ZBTB16 deficiency impairs endothelial cell function and induces premature aging

ZBTB16 plays a pivotal role in diverse cellular processes, including proliferation, differentiation, and inflammation^25^. Notably, ZBTB16 has been shown to restrict enhancer activity during hematopoietic progenitor aging^25^ and regulate autophagy^20^. However, its specific role in cardiac aging remained unexplored. To investigate its endothelial-specific functions, ZBTB16 was silenced in human umbilical vein endothelial cells (HUVECs) using siRNA (**Fig. 2A**). *ZBTB16* knockdown significantly impaired EC network formation, reduced VEGFA-induced migration, and inhibited angiogenic sprouting (**Fig. 2B-D**). Network formation was also inhibited in microvascular endothelial cells (**Supplementary data online Fig. 6**). Cell numbers decreased, while apoptosis and necrosis remained unaffected (**Fig. 2E-F**) suggesting a reduction of cellular proliferation. Importantly, *ZBTB16* silencing increased cellular senescence, as evidenced by elevated senescence-associated acidic β-galactosidase activity (**Fig. 2G**). These findings suggest that ZBTB16 plays a critical role in maintaining endothelial cell function and preventing senescence.

**Figure 2:**
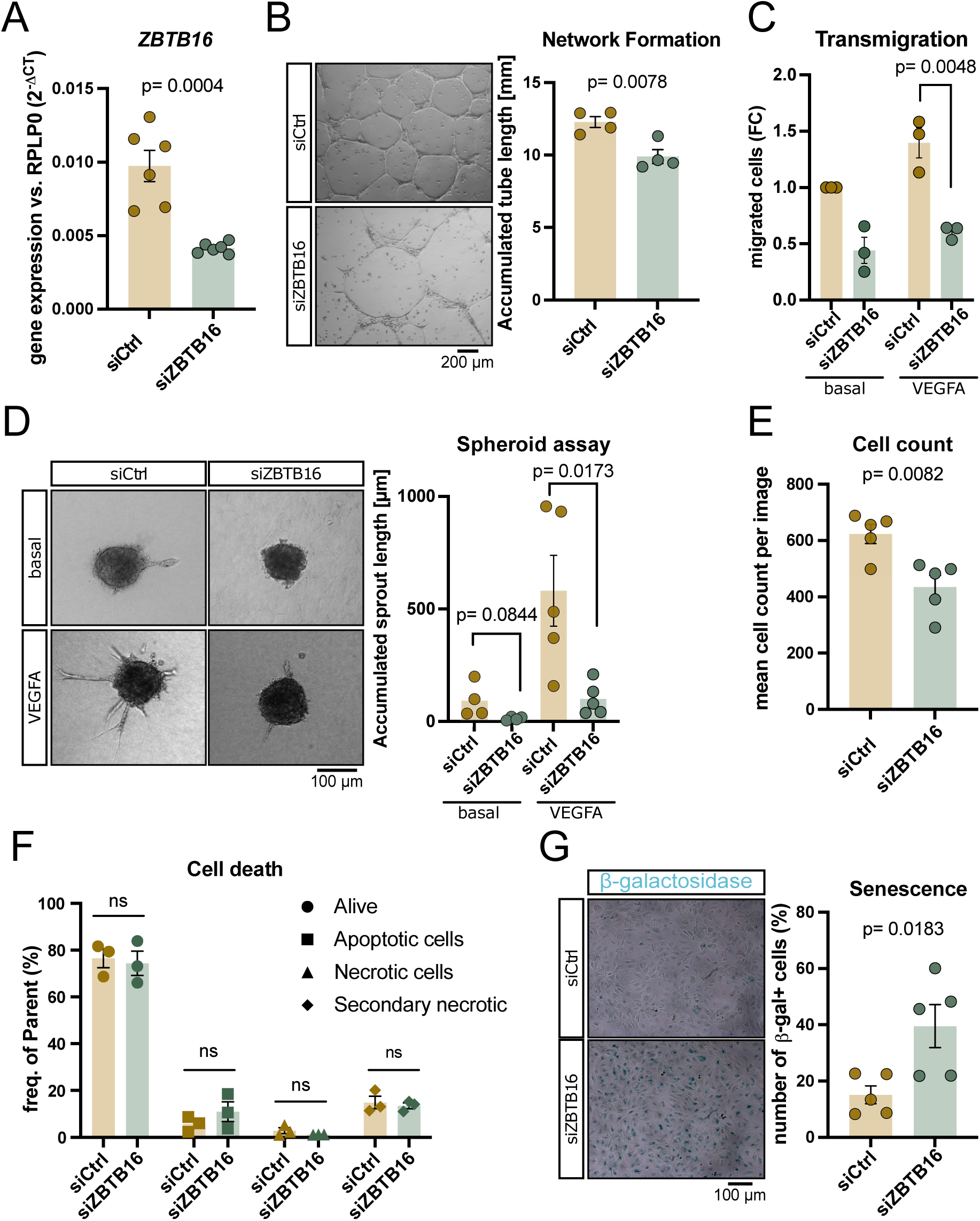
ZBTB16-silecing impairs endothelial function and induces cellular senescence. (**A**) siRNA mediated knockdown of ZBTB16 in human umbilical vein endothelial cells (HUVEC). Gene silencing was confirmed 72h after siRNA transfection (n=6). (**B**) Network formation assay of HUVEC transfected with siCtrl or siZBTB16 (n=4). (**C**) Quantification of Boyden Chamber transmigration assay using HUVEC after siCtrl Vs. siZBTB16 transfection. Data are normalized to siCtrl-treated cells without VEGFA. (**D**) Spheroid sprouting assay of HUVEC transfected with siCtrl or siZBTB16 followed by treatment with or without VEGFA (n=4 and n=5). Representative images are shown in the left panel. (**E**) Cell numbers were counted by visual assessment after 72h of knockdown (n=5). (**F**) Quantification of Annexin V and 7-AAD FACS. (n=3). (**G**) Representative images of senescence associated-β-galactosidase staining (senescence) at 72 h of siRNA treatment. Quantification is shown in the right panel (n=5). Data are shown as mean and error bar indicate the standard error of the mean. P-value was calculated by two-tailed Student’s t-test.

To explore ZBTB16’s role in vivo, heterozygous *Zbtb16* knockout mice (*Zbtb16+/-)* were studied, as homozygous knockout mice exhibited severe developmental abnormalities, including skeletal deformities and increased lethality (**Supplementary data online Table 1**). *Zbtb16+/-* mice, with reduced *ZBTB16* expression in endothelial cells (**Fig. 3A, Supplementary data online Fig. 7**), displayed premature cardiac aging phenotypes as early as 3-4 months. While body weight was not different between *Zbtb16+/+* and *Zbtb16+/-* mice (**Supplementary data online Fig. 8A**), echocardiographic analysis revealed increased E/É ratios, indicative of diastolic dysfunction, and elevated left ventricular mass (**Fig. 3B-C**), which are both hallmarks of aging-related cardiac dysfunction. Of note, diastolic dysfunction was more pronounced in female mice (**Supplementary data online Fig. 8B-D**). Systolic function as indicated by ejection fraction and blood pressure were unchanged (**Fig. 3D, Supplementary data online Fig. 8E**). Histological assessments confirmed age-associated changes, such as increased fibrosis and cardiomyocyte hypertrophy (**Fig. 3E-F**). Markers of cellular senescence were augmented in *Zbtb16*+/- hearts (**Fig. 3G, Supplementary data online Fig. 9**). Endothelial dysfunction, characterized by reduced capillary density and impaired outgrowth in an ex vivo aortic ring assay, was also observed (**Fig. 3H-I**). SnRNA-sequencing of *Zbtb16*+/- hearts further confirmed alterations in all major cardiovascular cell types (**Supplementary data online Fig. 10**). Particularly, capillary endothelial cells of *Zbtb16*^+/-^ mice showed an induction of genes associated with processes or GO terms such as “cardiac muscle hypertrophy” and “diabetic cardiomyopathy”, while GO terms such as “autophagy” and “DNA damage response” were down-regulated (**Supplementary data online Fig. 10**). In addition, we found increased processes associated with cardiac muscle hypertrophy or contraction in cardiomyocytes and alterations in gene expression patterns in fibroblasts of Zbtb16^+/-^ mice hearts (**Supplementary data online Fig. 11**).

**Figure 3:**
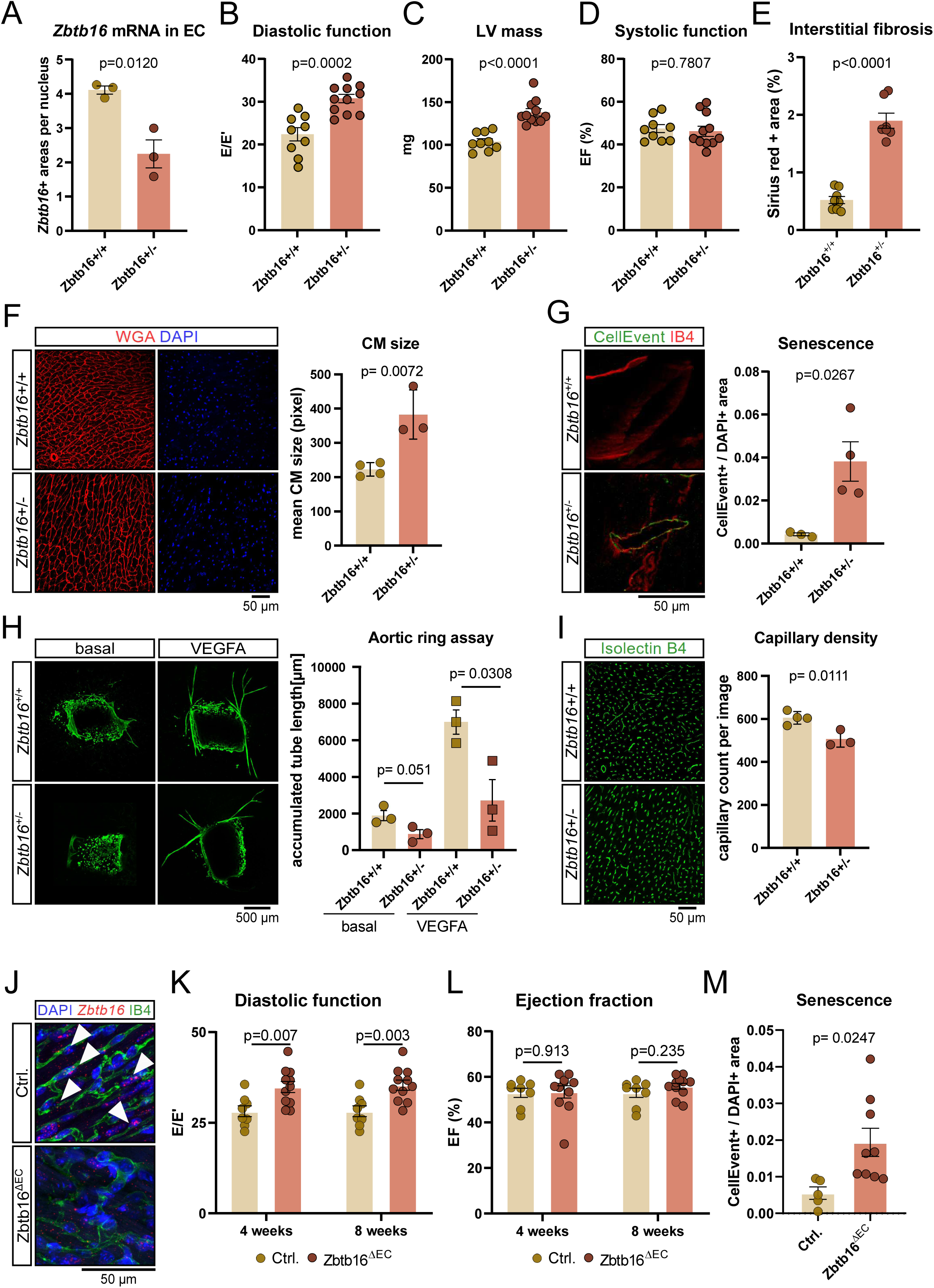
Heterozygous *Zbtb16*-deletion induces a pre-mature heart aging phenotype in young mice. (**A**) Quantification of *Zbtb16* positive endothelial cells in the heart of 3- to 4-month-old Zbtb16+/+ or Zbtb16+/- mouse hearts. *Zbtb16* mRNA was visualized by RNA-scope (n=3). Representative images are shown in Extended Data Fig. 5A. (**B-D**) Echocardiography in female (n=5 vs. n=5) and male (n=4 vs. n=6) Zbtb16+/+ versus Zbtb16+/- mice (3- to 4-month-olds). Data show diastolic function as E/E’ (B) and left ventricular mass in mg (C). Systolic function as ejection fraction is shown in panel D. (**E**) Quantification of Pico-Sirius Red stained areas (interstitial fibrosis by sparing vessels) in heart sections of 3- to 4-month-old *Zbtb16^+/+^*and *Zbtb16^+/-^* mice (n=8 vs. n=7). (**F**) Immunofluorescence images showing DAPI (blue) and wheat germ agglutinin (WGA) were used to measure cell size in cardiomyocytes (CM) *Zbtb16^+/+^* and *Zbtb16^+/-^* mice. Quantification of mean cardiomyocyte area is shown in the right panel (n=4 vs. n=3). (**G**) Senescence-associated β-galactosidase staining (CellEvent, green) of *Zbtb16^+/+^* and *Zbtb16^+/-^* mouse hearts (n=3 vs. n=4). Quantification of SA-β-galactosidase positive areas are shown in the right panel. (**H**) Ex vivo aortic ring assay of *Zbtb16^+/+^*and *Zbtb16^+/-^* mice (n=3). Left panel indicate basal conditions. Right panel was stimulated with VEGFA. Quantification is shown right. (**I**) Capillary density (Isolectin B4, IB4, green) quantification in images of hearts of *Zbtb16*^+/+^ and *Zbtb16*^+/-^ mice (n=3 vs. n=4). (**J-M**) Depletion of *Zbtb16* in endothelial cells in 3- to 4-month-old Cdh5-Cre; Zbtb16fl/fl (Zbtb16^DEC^ mice) or Cdh5-Cre-negative littermates (Ctrl.) which were treated with tamoxifen (8 weeks after start of treatment). (J) Representative images of *Zbtb16* mRNA (red) in capillaries (white arrow heads) as assessed by RNA-scope in heart sections after 8 weeks of tamoxifen injection. DAPI (blue) and IB4 (green) serve as counter stain. (K, L) Echocardiographic analysis of heart function (n=8 vs. n=11), 4 and 8 weeks after tamoxifen injection Shown are diastolic function as MV E/E’ (k) and systolic function as ejection fraction (L). (M) Histological analysis of senescence using the CellEvent staining kit (n=5 vs. n=8). Data are shown as mean and error bar indicate the standard error of the mean. P-value was calculated by two-tailed Student’s t-test.

To investigate the specific role of ZBTB16 in endothelial cells, we generated endothelial-specific knockout mice (Zbtb16^ΔEC^) by crossing Cdh5-ERT2Cre with *Zbtb16*^fl/fl^ mice (**Fig. 3J**). Four and eight weeks after tamoxifen induction, we observed premature aging phenotypes such as diastolic dysfunction (**Fig. 3K**) without alteration of systolic function (**Fig. 3L**). In line with the data from constitutive *Zbtb16*-deletion mice (**Fig. 3G**) and aged mouse hearts^8^, endothelial senescence was elevated in cardiac arteries from Zbtb16^ΔEC^ mice (**Fig. 3M**). These results highlight the essential role of endothelial Zbtb16 in protecting against premature cardiac aging.

### *Zbtb16*-deficiency causes endothelial paracrine dysfunctions

To further investigate ZBTB16’s regulatory role, bulk RNA sequencing was performed on ZBTB16-silenced endothelial cells. Transcriptomic analysis revealed that *ZBTB16* silencing increased inflammatory and fibrotic gene expression, but downregulated neuroprotective and angiogenic genes (**Fig. 4A**). Given the role of the endothelial niche in influencing surrounding cells through paracrine signaling, ZBTB16 loss may alter the endothelial secretome towards a pro-fibrotic and “inflammaging” phenotype, contributing to activation of neighboring fibroblasts or cardiomyocytes. Indeed, conditioned medium of *ZBTB16*-silenced endothelial cells activated human cardiac fibroblasts, inducing expression of α-smooth muscle actin and collagen-1A1 and stimulating fibroblast contraction (**Fig. 4B-F**). A three-dimensional cardiosphere model confirmed increased fibroblast activation and collagen deposition induced by *ZBTB16*-silenced endothelial cell derived conditioned medium (**Fig. 4G-I**). These data demonstrate that endothelial ZBTB16 loss promotes fibroblast activation via paracrine signaling.

**Figure 4:**
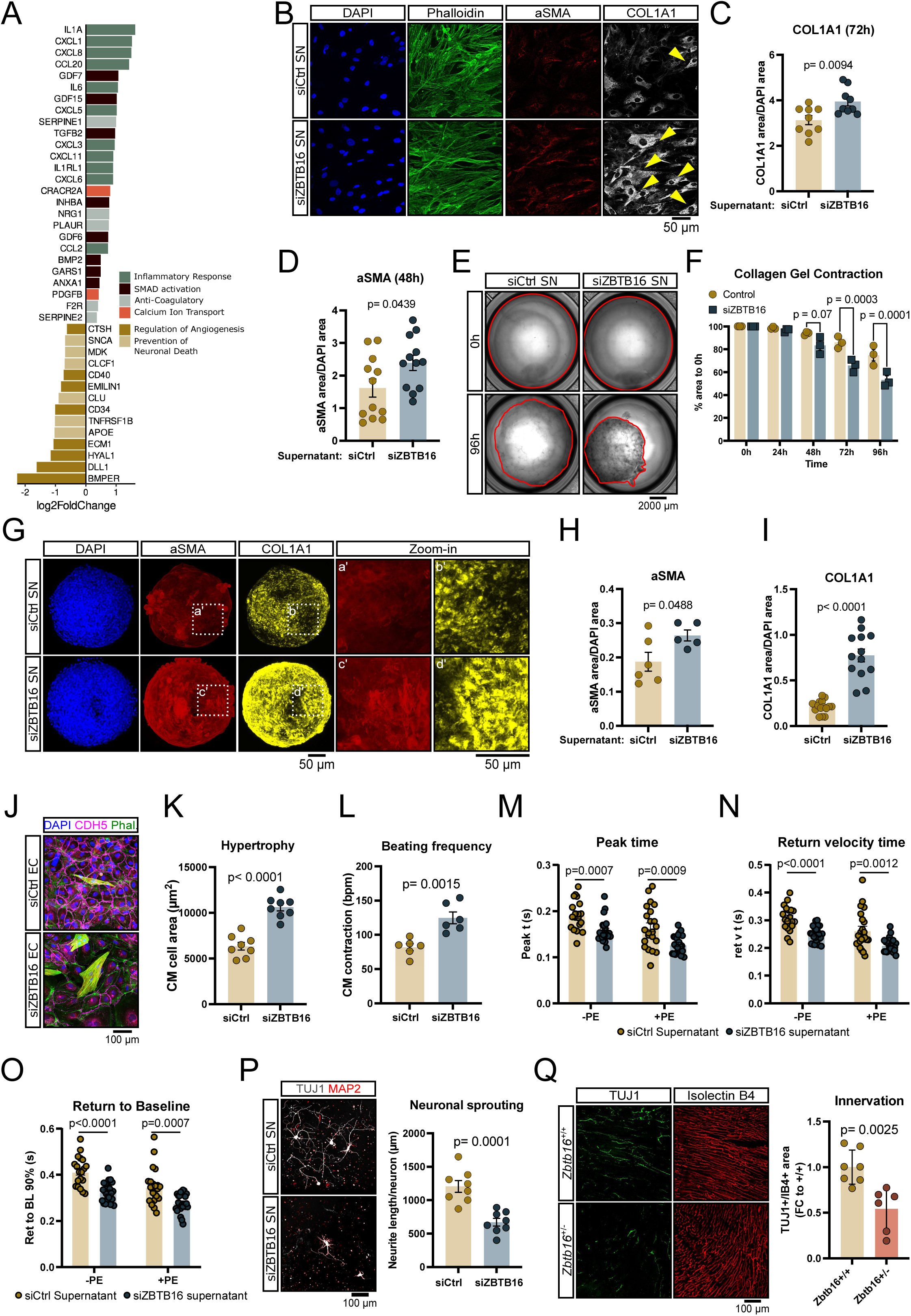
Conditioned media from *ZBTB16*-deficient HUVEC induces adverse changes in cardiac cells. (**A**) Bulk RNA sequencing of ZBTB16 knockdown cells and siRNA control cells (n=5). Graph shows log2 fold-change of genes associated with senescence (based on SenMayo (PMID: 35974106)). Genes were significant (Bonferroni adjusted p-value < 0.05). Positive log2 fold-change indicates higher expression in ZBTB16 knockdown cells. Genes were annotated according to their biological function. (**B-D**). Fibroblasts treated 72h with supernatants of HUVEC, which were transfected with siCtrl or siZBTB16 for 72h (n=10). Arrows indicate COL1A1 expression. Quantification is shown in c for COL1A1 (72h) and d for aSMA (48h). (**E-F**) Collagen gel contraction assay at baseline (upper panel) and after 96h (bottom panel) of fibroblast gels treated with supernatants from siCtrl (left) or siZBTB16 (right). Red line indicates gel boundaries. Quantification of relative decrease in gel area is shown in F. (**G-I**) Cardiac tissue mimetics (CTMs) containing primary rat cardiomyocytes, fibroblasts and endothelial cells were treated with supernatants of siCtrl or siZBTB16-transfected HUVEC for 14d. Representative immunohistochemical stainings are shown in G. Quantification of alpha smooth muscle actinin (aSMA) (H) and collagen COL1A1 (I). (**J-L**) siCtrl or siZBTB16-transfected HUVEC were directly co-cultured with neonatal rat cardiomyocytes (n=8) for 96h and stained for DAPI (blue), Phalloidin (green), sarcomeric-actinin (red) and VE-cadherin (magenta). Arrows indicate cardiomyocytes depicted by high sarcomeric actinin content. Cardiomyocyte hypertrophy quantified in panel k, and cardiomyocyte beating frequency is shown in L. (**M-O**) Neonatal rat cardiomyocytes were cultured in the supernatant of HUVEC after siCtrl or siZBTB16 transfection. Contraction (peak time) and relaxation (return velocity time, return to baseline 90%) were determined using IonOptix. Single cardiomyocytes were analyzed in the presence and absence of phenylephrine (-PE: n=19 vs. n=22; +PE: n=22 vs. n=23). (**P**) Primary mouse cortical neurons were treated with supernatants of siCtrl or siZBTB16-transfected HUVEC. Quantification is shown in the right panel. (**Q**) Innervation as assessed histologically by TUJ1 (green) normalized to IB4 (red) in hearts of *Zbtb16^+/+^* and *Zbtb16^+/-^*mice (n=6 vs. n=7). Quantification is shown in the right panel. Data are shown as mean and error bar indicate the standard error of the mean. P-value was calculated by two-tailed Student’s t-test.

In addition, we performed direct co-culture experiments of *ZBTB16*-silenced endothelial cells with isolated neonatal rat cardiomyocytes. Here, *ZBTB16*-silenced endothelial cells increased cardiomyocyte size and beating frequency, which are markers of cellular stress (**Fig. 4J-L**). In addition, single cardiomyocyte analyses revealed impaired cardiomyocyte contraction (**Fig. 4M**) and relaxation (**Fig. 4N-O**) in the presence of *ZBTB16*-repressed endothelial cell supernatant. Since endothelial cell aging was recently shown to reduce cardiac innervation^8^, we additionally determined the effect of conditioned medium from *ZBTB16*-silenced cells on neurite outgrowth in cultured neurons. We show that the conditioned medium reduces neurite outgrowth in vitro (**Fig. 4P**), a finding which is consistent with a reduced nerve density observed in the left ventricles of *Zbtb16*+/- mice (**Fig. 4Q**). These results suggest that ZBTB16 downregulation in endothelial cells contributes to impaired neurovascular and endothelial-cardiomyocyte interactions during aging.

To identify down-stream targets mediating the effects of ZBTB16, we performed ZBTB16 chromatin-immunoprecipitation studies (**Supplementary data online Fig. 12A-B, Supplementary data online Table 2**). We identified *NRIP1* (Nuclear receptor-interacting protein 1) as a potential direct target of ZBTB16 (**Supplementary data online Fig. 12C-D**). Since *Nrip1* expression was elevated in aged mouse cardiac endothelial cells (**Supplementary data online Fig. 13A**), we hypothesize that ZBTB16 may repress *Nrip1*. Indeed, silencing of *ZBTB16* in endothelial cells augments *NRIP1* expression (**Supplementary data online Fig. 13B**). To address if NRIP1 may contribute to the ZBTB16 regulated paracrine activation of fibroblasts, we co-silenced *NRIP1* with *ZBTB16* (**Supplementary data online Fig. 13C**). Silencing NRIP1 indeed blunted siZBTB16-induced fibroblast activation and collagen-1A1 expression (**Supplementary data online Fig. 13D-E**), further implicating NRIP1 in ZBTB16-mediated paracrine signaling.

In addition, we further assess the downstream targets of ZBTB16 controlling endothelial-nerve cross-talks. Bulk ATAC- and RNA-sequencing of siZBTB16-versus control-transfected HUVEC showed GO terms associated with ’axons’ or ’neuron projection guidance’ as the top down-regulated pathway (**Supplementary** Fig. 14A-B). We particularly found ephrin family members to be included in these GO terms. Ephrins such as EFN-4 are known to act as axon guidance cue in *C. elegans*^26^. Human and mouse orthologs for EFN-4 are EFNB1 and EFNB2. Interestingly, both genes and other members of the ephrin family are down-regulated in HUVEC upon ZBTB16 silencing (**Supplementary data online Fig. 14C**). To test the functional relevance, we silenced the highest expressed ephrin, EFNB2, by siRNA in HUVEC and transferred the supernatant of these cells to cortical neurons. Indeed, supernatants from siEFNB2-transfected HUVEC significantly impaired axon sprouting compared to control transfected HUVEC (**Supplementary data online Fig. 14D**). These data suggest that ZBTB16 might control axon sprouting by regulating EFNB2 expression.

### Therapeutic potential of ZBTB16 overexpression

Finally, the therapeutic potential of ZBTB16 in restoring a healthy phenotype in senescent cells was tested. Lentiviral overexpression of *ZBTB16* in long term cultured senescent endothelial cells in vitro improved angiogenic capacity, reduced senescence, and mitigated fibroblast activation induced by the aged endothelial secretome (**Fig. 5A-C**). To further explore the therapeutic potential in vivo, we overexpressed *Zbtb16* in murine endothelial cells using adeno-associated viral vectors, which were coated with an endothelial-targeting peptide linked to PAMAM and express the target gene from an endoglin promotor to enrich for endothelial tropism^8,22^ as evidenced by RT-qPCR on isolated liver EC and by RNA-scope on heart sections (**Fig. 5D, Supplementary data online Fig. 15**). Endothelial-specific overexpression of *Zbtb16* indeed prevented the further decline in age-associated diastolic dysfunction (**Fig. 5E, Supplementary data online Fig. 16A**) and elevated ejection fraction in aged mice, as well (**Fig. 5F**). Senescence tends to be reduced (p=0.09) (**Fig. 5G**). While interstitial fibrosis revealed a trend wise reduction by 15 % (**Supplementary data online Fig. 16B**), perivascular fibrosis was significantly reduced to levels of the 18 months old mice demonstrating that *Zbtb16* overexpression prevents the further progression of fibrosis in aged mice (**Fig. 5H**). These findings highlight ZBTB16 as a potential therapeutic target for mitigating endothelial dysfunction and cardiac aging.

**Figure 5:**
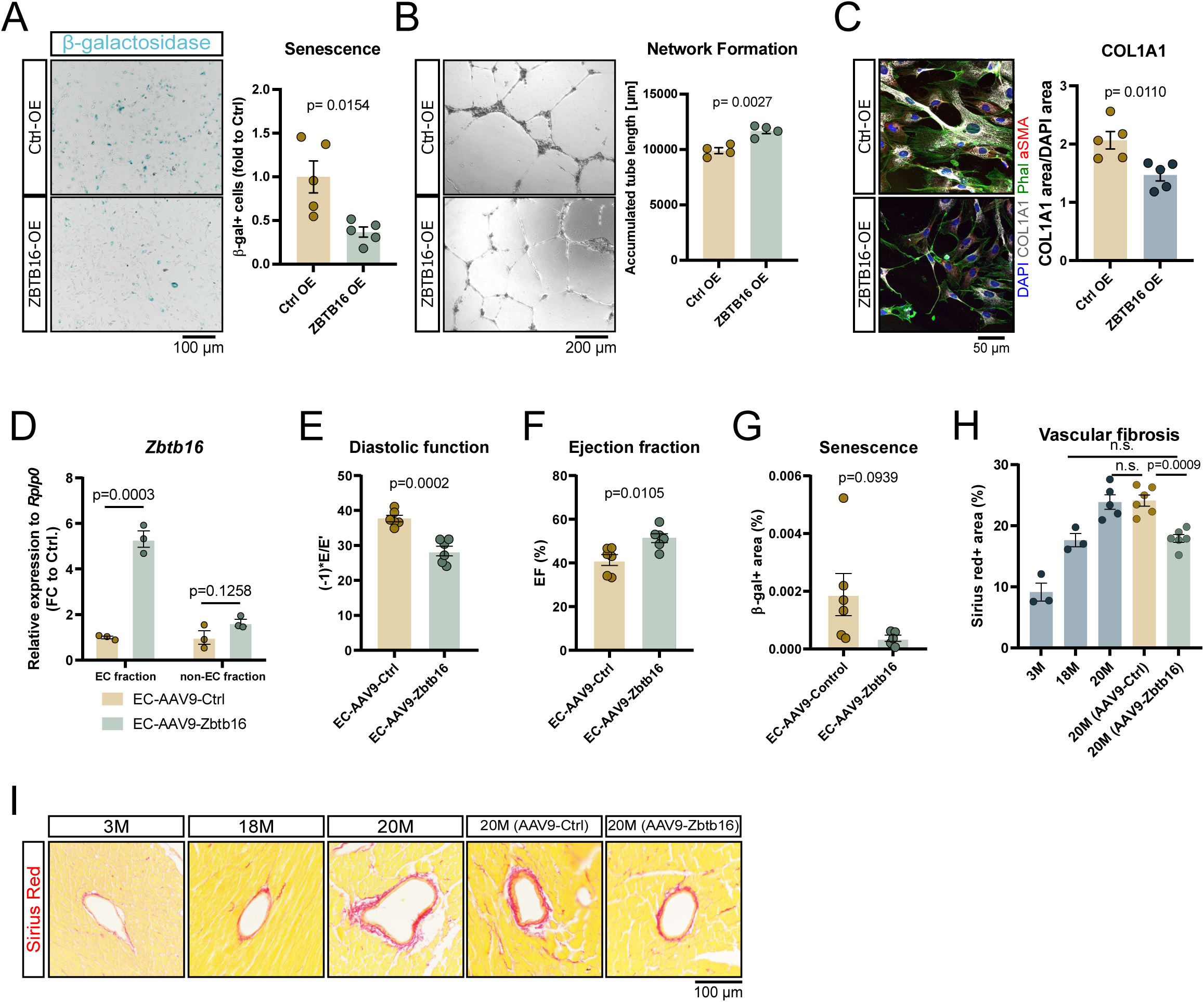
*ZBTB16*-overexpression in senescent endothelial cells rescues endothelial dysfunction. (**A-C**) *ZBTB16* was overexpressed by lentiviral vectors in long-term passaged HUVEC (>P9) for >8 days. (A) Staining of HUVEC for acidic-β-galactosidase (n=5). (B) *ZBTB16* was overexpressed in long-term passaged senescent HUVEC (>P12) prior to performing a network formation assay (n=4). (C) *ZBTB16* was overexpressed in long-term passaged HUVEC (>P12). Supernatants were collected and transferred to human cardiac fibroblasts (n=5). After 72h, fibroblasts were stained for COL1A1 (grey), DAPI (blue), aSMA (red) and Phalloidin (green). Scale bar indicates 50 µm. (**D-I**) Overexpression of Zbtb16 in endothelial cells by targeted AAV9 vectors improves cardiac function in 18-month-old mice. (D) *Zbtb16* expression in liver endothelial cells 4 weeks after AAV9 treatment. (E) Diastolic heart function and F ejection fraction 8 weeks after AAV9 treatment. Acidic β-galactosidase (G) of cardiac sections 8 weeks after AAV9 treatment. (H, I) Vascular fibrosis as assesses by Sirius red staining on heart section from 3-month-old (n=3), 18-month-old (n=3), 20-month-old (n=5) and 20-month-old mice after two months of ZBTB16-AAV9 or control treatment (n=6). Data are shown as mean and error bar indicate the standard error of the mean. P-value was calculated by two-tailed Student’s t-test or using One-way ANOVA with a post-hoc Tukey’s test (H).

## DISCUSSION

Our study reveals a critical role of ZBTB16 in preserving endothelial function and mitigating cardiac aging. Constitutive or endothelial-specific ZBTB16 deficiency induces premature aging through disrupted paracrine signaling, promoting fibrosis and hypertrophy. This supports the vascular niche’s key paracrine role in tissue homeostasis. Aging alters the vascular niche, impairing microcirculatory function, reducing oxygen and nutrient supply, and promoting inflammation. Endothelial senescence triggers the release of SASPs. These include cytokines and growth factors that affect neighboring cells. Loss of ZBTB16 exacerbates these effects, contributing to endothelial dysfunction. In vitro studies confirm that supernatants from ZBTB16-deficient endothelial cells exhibit pro-fibrotic and pro-hypertrophic activity, implicating paracrine dysfunction in premature aging. Of note, overexpression of endothelial ZBTB16 reverses these impairments, reverting endothelial dysfunction and age-related cardiac diastolic dysfunction. These findings highlight ZBTB16 as a promising therapeutic target for combating vascular aging.

In contrast to our data, ZBTB16 has been shown by others to attenuate fibrosis and cardiac hypertrophy^27^. However, this study used Zbtb16-/- mice, which exhibited reduced body weight and, in our hands, showed embryonic lethality. Our findings align with previous reports demonstrating that biallelic loss of Zbtb16 is associated with congenital heart defects^28^, raising concerns about the validity of using the few homozygous mice that survive beyond the embryonic stage. Another group^29,30^ used spontaneously hypertensive rats and demonstrated that heterozygote Zbtb16 rats showed improved metabolic and cardiac traits. However, also in this study the authors reported that the targeted allele is semi-lethal and report altered body weights, which we did not observe in our mice cohorts.

Therefore, the protective effects seen in these studies may reflect compensatory pathways in survivor mice.

Our data provide additional insights into the transcriptional and epigenetic control of cardiac endothelial cell aging. While we focused on the assessment of ZBTB16 functions, which was repressed during aging, our snATAC-seq data suggest that additional genes involved in the control of neuro-vascular cross talks, angiogenesis, barrier function and transporters are regulated. Since epigenetic alterations are considered long lasting, these effects on gene expression may contribute to chronification of the aging signatures. The mechanisms underlying the epigenetic control of ZBTB16 and the other regulated gene loci needs to be further studied.

Mimicking *Zbtb16* repression during aging by haploinsufficient depletion of the gene, induced profound alterations of gene expression. ZBTB16 is well known to act as a transcriptional regulator by binding to specific DNA sequences via its zinc finger domains^16^ and additionally recruits co-repressor complexes via its BTB domain^31,32^. Further studies pointed to a role as probable substrate-recognition component of an E3 ubiquitin-protein ligase complex, which mediates the ubiquitination and subsequent proteasomal degradation of target proteins^33,34^. The complex mechanisms of action are mirrored by several functional effects in vitro. ZBTB16 has been proposed a pro-proliferative^19^ and autophagy-inducing function^20^. However, another study showed an inhibition of angiogenesis by ZBTB16 overexpression in endothelial cells, which seems not to express Zbtb16 at baseline^35^, which is not consistent with our findings and publicly available sequencing results. Our data suggest that ZBTB16 binds to the promoters of several genes and repressed NRP1. NRIP1 has been identified as a critical downstream target, through which loss of ZBTB16 exerts its effects on fibroblasts. While little is known regarding the function of NRP1 in endothelial cells and the vasculature, it is known for its effect in metabolic and other diseases. NRIP1 is augmented in preclampsia^36^, and aggravates pulmonary microvascular injury during inflammation^37,38^. It additionally has broader metabolic effects, where NRIP1 deficiency improves cardiac metabolism^39^, augments fat utilization^40^ and can extend female longevity^40^. As ZBTB16 is induced by energy deficit conditions like fasting, glucopenia, and cold exposure in the brain^41^, it may be worth studying the impact of metabolic interventions, which are known to interfere with aging^42,43^, on cardiac endothelial cell ZBTB16 expression. These findings may further link ZBTB16 to broader metabolic and inflammatory pathways in aging. Exploring therapeutic strategies to induce ZBTB16 or interfere with its down-stream signaling pathways may open new avenues for combating cardiovascular aging.

Our study additionally shows that Zbtb16 regulates the expression of ephrin family members. Ephrins have multiple effects in cell-cell-communication. Particularly, we showed that silencing of EFNB2 inhibits endothelial-nerve interactions. Moreover, the effect of Zbtb16 on Ephrin expression may additionally impact endothelial-cardiomyocytes interaction given that previous studies demonstrated that silencing of EFNB2 in endothelial cells induces cardiomyocyte hypertrophy in coculture studies^23^. Therefore, it is conceivable that inhibition of ZBTB16 in old heart endothelial cells reduces EFNB2, thereby decreasing axons and increasing cardiomyocyte hypertrophy.

### Limitation of the study

While our study demonstrates that endothelial overexpression of ZBTB16 can prevent cardiac aging, significant translational hurdles remain before this finding can be developed into a viable therapeutic strategy. The use of viral-based delivery methods in the present study, though effective experimentally, presents limitations for clinical application due to concerns regarding safety and immunogenicity. As an alternative, nanoparticle-based delivery systems may offer a more clinically feasible approach. Nonetheless, the complex and multifaceted role of ZBTB16, along with the ongoing challenges in achieving safe, efficient, and sustained endothelial-specific delivery in humans, underscores the need for continued research to bridge this translational gap. In addition, the mechanisms regulating *Zbtb16* expression, as well as the pathways through which it influences cardiac aging, remain to be fully elucidated. While our in vitro experiments indicate that *Zbtb16* silencing activates cardiac fibroblasts via NRIP1, it is likely that *Zbtb16* exerts additional effects on other cardiac cell types and may even have systemic roles by modulating other vascular beds. Furthermore, the limitations of the cellular models, such as the use of HUVEC and neonatal rat cardiomyocytes, must be acknowledged, and the in vivo function of NRIP1 in this setting has yet to be explored.

In conclusion, this study identifies ZBTB16 as a key endothelial regulator that preserves cardiac function during aging by maintaining vascular niche homeostasis and limiting pro-fibrotic signaling. Loss of ZBTB16 accelerates endothelial senescence and promotes diastolic dysfunction, while its overexpression reverses these effects in aged mice. Further work is needed to explore the use of ZBTB16 as therapeutic target for preventing age-related cardiac dysfunction.

## Acknowledgements

We thank Alisa Debes for experimental support and Dennis Hecker as well as Marcel Schulz for bioinformatics support.

## Data and materials availability

All data are available in the manuscript or the supplementary materials. Material is available upon request from the authors indicated in the material section. Sequencing data will be available on ArrayExpress upon accession number XXXX. Bulk ATAC sequencing data can be accessed via the accession number GSE303574.

## Contributions

Conceptualization: J.U.G.W., S.D., T.Br., RPB

Wet lab experiments: K.A.S., S-F.G., V.E.L., V.L., W.A., J. P., H. K., A.F., K.S., M.Y., S.G., T.S., M-D.P., J.K., PFM

Bioinformatic analysis: D.R.M., M. R. J., L.T., D.J., M.Y., S.G.

Provision of material: O.M., S.H., C.K., T.B., H. S., M. O.

Funding acquisition: J.U.G.W., S.D.

Supervision: J.U.G.W. and S.D.

Writing – original draft: K.S., J.U.G.W., S.D.

Writing – review and editing: A.M.Z., D.R.M.; all authors have read and approved the current manuscript.

## Ethics declarations

Competing interests: K.S., A.Z., S.D., J.U.G.W have applied for a patent

## Supplementary data online

Supplementary data online Fig. 1-16

Supplementary data online Table 1-2

Extended Material & Methods

## Supplementary material and methods

### Laboratory animals and in vivo experiments

C57Bl/6J wildtype mice were purchased from Janvier (Le Genest SaintIsle, France) and from Charles River (Sulzfeld, Germany). Homozygosity of these inbred mice was controlled by Janvier and Charles River using exome sequencing. Zbtb16-deletion mice (Strain ID: 066990-UCD) were purchased from UC Davis MMRRC (Los Angelos, CA, USA). Zbtb16-fl/fl mice were purchased from GemPharma Tech (strain number: T016043). All animal experiments have been executed in accordance to the guidelines from Directive 2010/63/EU of the European Parliament on the protection of animals used for scientific purposes and were approved by the local authorities by the state of Hesse (Regierungspräsidium Darmstadt). Experiments were conducted as follows: mice were anaesthetized with 2-2.5% of isoflurane and echocardiography was performed to monitor heart function using the Vevo 3100 echocardiography system with the Vevo LAB software (Fujifilm VisualSonics). Systolic function (ejection fraction) and diastolic function (expressed as E/E’ * (-1)) were measured. At the end of the experiment, mice were anaesthetized with 2-2.5% of isoflurane and euthanized via cervical dislocation.

### Tail-Cuff Measurement

Blood pressure measurement using the tail-cuff technique (tail plethysmography). A standard device from Vistech (BP2000) was used. Blood pressure was automatically determined using a small cuff and a laser sensor near the base of the tail. To minimize stress-induced changes in blood pressure, the mice were acclimated to the procedure on several consecutive days before the actual experiment began. To ensure stable measurements, the mice were placed individually in small metal boxes during the measurement, in which the tail is guided out through a hole. Since the mouse tail is thermoregulatory active, the box is warmed to 34°C to ensure good blood flow to the tails, comparable between groups. The tail-cuff measurement is carried out at most once per day, with a maximum of 7 days for acclimation and 3 times for blood pressure measurement.

### AAV9-mediated Zbtb16 induction

To induce endothelial-targeted *Zbtb16* expression, AAV9 vectors encoding the murine *Zbtb16* or fLuc2 under the control of the *endoglin* promotor were generated. Viral particles were coated with a G2-PAMAM linked to an endothelial-targeted peptide (PAMAM-G2^CNN^) which were generated as previously described^1,2^ and referred to as EC-AAV9-Zbtb16. Viral particles were diluted in OptiMEM GlutaMAX^TM^ (51985026, Gibco) and coated with PAMAM G2CNN (1.8 µg of PAMAM-G2^CNN^ to coat 2×10^12^ viral particles) for 30 minutes at room temperature, prior to injection. 2×10^12^ AAV9-Zbtb16 were i.v. injected to 18 months old C57Bl/6J mice. AAV9 encoding for fLuc2 served as negative control. Four or eight weeks after injection, mice were euthanized for histological and transcriptomic assessments. The animal experiment has been conducted as approved by the state of Hessen.

### Aortic ring assay

After mice were euthanized, hearts were flushed with ice cold Hank’s buffered saline solution (HBSS). The aorta was dissected sectioned into at least 8 rings. Each ring was embedded in 50µl rat tail collagen type 1(354236, Corning) in 1x medium M119 (M0650, Sigma-Aldrich), 8 mM NaHCO_3_ (HN01.2, Carl Roth GmbH) and 10 mM NaOH (6771.1 Carl Roth GmbH) in a 96 well plate. Aortic rings were incubated at 37°C in order to add 150 µL DMEM/F-12 (10565018, Gibco) supplemented with 2.5% FBS (4133, Invitrogen) per well.

The aortic rings were cultured in presence or absence of 60 ng/mL of human recombinant VEGFA (SRP4363-10UG, Sigma Aldrich). After 7 days in culture, aortic rings were fixed with 4% PFA (28908, Thermo Scientific) for 30 minutes and permeabilized with 0.25% Triton-X containing PBS for 15 minutes. After blocking with Dako protein blocking buffer (X0909, DAKO), aortic rings incubated overnight in PBLEC buffer (50 µl 1M MgCl_2_, 50 µl 1M CaCl_2_, 5 µl 1M MnCl_2_, 5 ml 10% Triton-X-100, in 50 ml PBS) containing biotinylated Isolectin B4(VEC-B-1205, Biozol) in a concentration of 1:100. Aortic rings were washed three times with PBS and stained with streptavidin conjugated Alexa 488 (S32354, Invitrogen) for 3 hours at room temperature and washed with PBS three times, with 15 minutes each wash. The stained rings were imaged using the Leica DMi8 confocal Stellaris microscope and analysed with the LASX software.

### RNA in situ hybridization (RNA-scope)

Murine hearts were dissected after perfusion with PBS and 4%PFA. Dissected hearts were fixed in 4% PFA for 24 hours at 4°C followed by an ascending sucrose gradient (10, 20, and 30% sucrose in PBS) for 24 hours at 4°C each. The hearts were embedded in optimal cutting temperature (OCT) medium and stored at -80°C for 24 hours or until sectioning. Tissue sections of 15 µm were obtained from cryostat (CM3050, Leica) and stored at -20°C. RNA in situ hybridization was performed for *Zbtb16* using a probe (437171-C3), Protease (III), and TSA Fluorophores (PN 323273), in RNA-scope Multiplex Fluorescent Reagent Kit v.2 (Advanced Cell Diagnostics, 323270), according to the manufacturer’s protocol. For antibody staining, the samples were incubated in a blocking buffer containing 3% bovine serum albumin (BSA, A7030, Sigma-Aldrich), 0.1% Triton X-100, and 5% donkey serum (ab7475, Abcam) for an hour at room temperature, following RNA in situ. Isolectin B4 and respective antibodies were diluted in blocking buffer and incubated overnight at 4°C in a humid chamber. The next day, after three washes with PBS for 5 minutes each, secondary antibody and DAPI were added in 5%BSA in PBS and incubated at room temperature for one hour, followed by three washes in PBS for 5 minutes. Slides were mounted in Fluoromont-GTM mounting media. Fluorescent images were acquired using Leica Stellaris confocal microscope. Quantification of RNA scope was performed in Image J according to the guidelines provided by Advanced Cell Diagnostics

### Immunolabeling methods

Paraffin sectioned hearts incubated at 60°C for 30 minutes. Sections were deparaffinized and rehydrated using Xylene (two times for 10 minutes) and a series of ethanol gradients (100%, 95%, 80%, 70% and 50% ethanol) (5 minutes each) respectively. Afterwards, sections were washed with water for 5 minutes and were incubated in boiling 0.01M citrate buffer, pH 6.0 for 90 to 120 seconds. Slides were blocked with PBS containing 0.1% Triton-X100, 3% BSA (A7030-10G, Merck), and 5% donkey serum (ab7475, Abcam). The sections incubated with the blocking solution that contained the primary antibodies overnight at 4°C and sections were washed three times in PBS for 5 minutes each wash and incubated with the corresponding secondary antibodies and DAPI (6335.1, Carl Roth GmbH & Co.KG) diluted in PBS containing 0.1% Triton X-100 for one hour at room temperature. The sections were washed with PBS three times for 5 minutes each and mounted with Fluoromount-G^TM^ (00-4958-02, Invitrogen). Images were taken using the Leica DMi8 confocal Stellaris microscope and the LASX software.

For cryo-preserved sections, hearts were flushed with cold HBSS were fixed overnight at 4°C in 4 %PFA in PBS (28908, ThermoFisher Scientific). The fixed hearts were washed with PBS three times, 10 minutes each wash. Hearts were then subjected to three consecutive overnight incubations at 4°C with increasing concentrations of sucrose (10%, 20%, 30%; S0389, Sigma-Aldrich) in PBS and the tissues were embedded in 15% sucrose with 8% gelatin (G1890, Sigma-Aldrich), and 1% polyvinylpyrrolidone (P5288, Sigma-Aldrich) in PBS. The embedded tissues were allowed to solidify and stored at -80°C. Hearts were sectioned using a cryostat (Leica CM3050 S) into 50 µm sections, mounted on adhesive glass slides (10149870, ThermoFischer Scientific) and stored at -20°C.

To stain cryopreserved sections, sections were allowed to acclimatize to room temperature for 5 minutes and were re-hydrated with PBS twice, 5 minutes each. Sections were permeabilized with PBS containing 0.3% Triton X-100 thrice, 10 minutes each and blocked with 0.1% Triton X-100, 3% BSA (A7030-10G, Merck) and 5% donkey serum (ab7475, Abcam) in PBS for 1 hour at room temperature. After blocking, sections incubated with primary antibodies (diluted in blocking solution), overnight at 4°C. Slides were washed with PBS three times, with 5 minutes each wash. After one additional step in 0.1% Triton X-100 in PBS, sections were incubated with corresponding secondary antibody in PBS containing 5% BSA (A7030-10G, Merck) for 1 hour at room temperature. Slides were then washed with PBS for 5 minutes, three times and mounted with with Fluoromount-G^TM^ (00-4958-02, Invitrogen). Images were taken using the Leica DMi8 confocal Stellaris microscope and the LASX software.

To stain cultured cells, cells were fixated with 4% PFA for 10 minutes at room temperature and washed with PBS containing 0.1% Triton-X for 10 minutes. Cells were blocked using 2% donkey serum (ab7475, Abcam), 1% BSA (A7030-10G, Merck) and 0.1% Triton-X in PBS for 1 hour and incubated with the primary antibodies overnight at 4°C. The cells were washed with PBS, three times for 5 minutes each wash and incubated with the corresponding secondary antibody for 1 hour at room temperature, followed by PBS wash for three times, 5 minutes each wash. The cells were then mounted with Fluoromount-G^TM^ (00-4958-02, Invitrogen) and imaged with Leica DMi8 confocal Stellaris microscope and the LASX software. Images were analyzed using software Volocity 7 by Quorum Technologies Inc. and normalized to IB4 or DAPI area.

Protein expression was determined via western blotting as previously described^3^. Hearts from C57Bl/6J wildtype mice (3- and 22-month-old) were homogenized and lysed in RIPA buffer (89900, Thermo Scientific) supplemented with 1x Protease inhibitor (cOmplete Mini, 11836170001, Roche), 1x Phosphatase Inhibitor (PhosSTOP, 04906837001, Roche). Membranes were incubated overnight at 4°C with primary antibodies (Rabbit anti-ZBTB16 (1:2000, A00817-1, Boster Biological Technology), Rabbit anti-GAPDH (1:1000, 2118S, Cell Signalling), followed by incubation with a secondary antibody (Donkey to Rabbit IgG (HRP), ab6802, Abcam).

### Antibodies used for this project

Primary antibodies: Rabbit anti-ZBTB16 (1:100, PA5-112862, Invitrogen), Mouse anti-ZBTB16 (1:100, sc-28319, Santa Cruz Biotechnology), Rabbit anti-COL1A1 (1:100, 72026S, CST), Rabbit anti-TUJ1 (1:100, Abcam, AB18207), GSL 1-Isolectin B4 (IB4) (1:25, Biozol, VEC-B-1205), Chicken anti-MAP2 (1:100, Abcam, ab5392), Rabbit anti-CD68 (CST, 97778), Wheat Germ Agglutinin (WGA), Alexa Fluor™ 555 Conjugate (1:100, W32464, Invitrogen) and Mouse anti-α-smooth muscle actin - Cy3™ (αSMA, 1:200, C6198-2ML, Sigma-Aldrich), Mouse anti-Actin (α-Sarcomeric) antibody (A2172-100UL, Merck), Mouse anti-α-actinin (1:200, A7811, Merck), Rabbit anti-CDH5 (1:200, 2500S, Cell Signaling), Rabbit anti-GAPDH (clone: 14C10, 1:5000, 2118S, CST).

Secondary antibodies: Alexa Fluor^TM^ 488 Phalloidin (1:100, Invitrogen, A12379), Streptavidin Alexa Fluor™ 488 conjugate (1:100, Invitrogen, S32354), Donkey anti-rabbit antibody Alexa Fluor 488 (1:200, Invitrogen, A-21206), Donkey anti-Rabbit Secondary Antibody Alexa Fluor 555 (1:200, Invitrogen, A-31572), Donkey anti-Chicken Alexa Fluor 488 (1:200, Dianova, 703-545-155), Donkey anti-Rabbit ECL IgG, HRP-linked (NA934-1ML, Amersham).

### Acidic Beta galactosidase staining

Senescence-associated β-galactosidase was carried out on cryopreserved sections using the senescence β-galactosidase staining kit (9860, CST) and the CellEvent™ Senescence Green Detection Kit (C10850, ThermoFisher) according to manufacturer’s instruction. The β-galactosidase positive areas were quantified using ImageJ or Volocity 7 (Quorum Technologies Inc.).

### Picrosirius red staining

Fibrosis was visualized using picrosirius red staining on paraffin sections. Sections were deparaffinized using wash with Xylene, twice for 10 minutes each wash and rehydrated using series of ethanol gradients (100%, 95%, 80%, 70% and 50% ethanol; 5 minutes each). After 5 minutes in water, sections incubated with 0.1% picrosirius red solution (0.5g sirius red in 500 ml of saturated aqueous picric acid) for 1 hour and washed twice with acidified water. The stained slides were dehydrated using 100% ethanol, cleared with xylene and mounted using Pertex mounting medium (R0080, Histoline).

### Cell culture

Human umbilical cord vein endothelial cells (HUVEC) were purchased from PromoCell (C-122203) and cultured in endothelial basal medium (EBM, CC-3121, Lonza) supplemented with 10% fetal bovine serum (10100147, Gibco). Cells were derived from mixed sexes.

*ZBTB16* repression was mediated by using the On-Target plus SMART pool siRNAs against human *ZBTB16* (L-018719-00-0005, Dharmacon / Horizon). AllStars Negative Control siRNA (1027280, Qiagen) served as negative control. Both siRNA pools (40 nM) were transfected in OptiMEM (31985070, Gibco) and Lipofectamin RNAiMAX (13778100, Invitrogen). To repress *NRIP1* expression the On-Target plus SMART pool siRNAs against human *NRIP1* (L-006686-00-0005, Dharmacon / Horizon) was used. The repression of EFNB2 was performed as previously described^4^.

Primary mouse cortical neurons (A15585, Gibco) were purchased and cultured according to manufacturer’s instruction in Neurobasal medium (21103049, Gibco) containing GlutaMAX-I (35050, Gibco) and B-27 supplement (17504, Gibco). Neurons were treated with conditioned HUVEC medium (50:50, Neurobasal^TM^ medium: conditioned medium).

Human cardiac Fibroblasts (HCF-c) primary cells (12375) were purchased from Promocell and cultured according to manufacturer’s instruction in fibroblast basal medium 3 (C-23230, Promocell) containing the recommended supplement mix (c-39350, Promocell). Fibroblasts were treated with TGF beta (302-B2-10, Biotechne) or conditioned HUVEC media for 48 or 72 hours.

Fibroblast activation was determined by collagen contraction assays as previously described^5^.

Neonatal rat cardiomyocytes were isolated and maintained as previously described^4^. Cardiomyocytes were seeded on a confluent HUVEC layer that has been transfected with siRNA pools against *ZBTB16* or non-targeting control. The detailed experimental procedure was previously described^4^.

### Lentivirus production and transduction

Plasmids containing the coding sequence of *ZBTB16* with HA tag or stuffer control under spleen focus forming virus (SFFW) promoter were purchased from vector builder. These plasmids were designed to contain GFP for transduction efficiency verification and puromycin resistance for the selection of transduced cells. The Lentivirus production process involved co-transfection of HEK293FT cells with pMD2.G, pCMVΔR8.91, and transfer plasmids as described before^6^. Transfections were performed with Genejuice transfection reagent (Millipore, 70967), the media was refreshed after 24 hours, and the virus was collected at 48 and 72 hours post-transfection. According to the manufacturer protocol, the collected virus was concentrated with the lentiX concentrator. Pooled HUVECs were transduced at passage 1, the media was refreshed after 24 hours and the transduced cells were allowed to expand another 48 hours. The percentage of GFP-positive cells assessed the efficiency of transduction, and overexpression of *Zbtb16* was confirmed by RT-qPCR.

### Liver endothelial cell isolation

Endothelial cells were isolated from murine liver as previously described^3^.

### RNA isolation

The miRNeasy Mini kit (217004, Qiagen) in comination with an on-column DNase I digestion (79254, Qiagen) was used for RNA isolation in accordance to the manufacturer’s instruction. The RNA concentration was determined by measuring absorption at 260 nm and 280 nm with the NanoDrop ND 2000-spectrophotometer (PeqLab).

### cDNA synthesis and quantitative PCR

Reverse transcription was performed using reverse transcriptase M-MLV (28025013, ThermoFisher Scientific) from 500 ng of RNA and assessed using the SYBR™ Green PCR Master Mix (4385617, Applied Biosystems). The primers were designed and purchased from Merck:

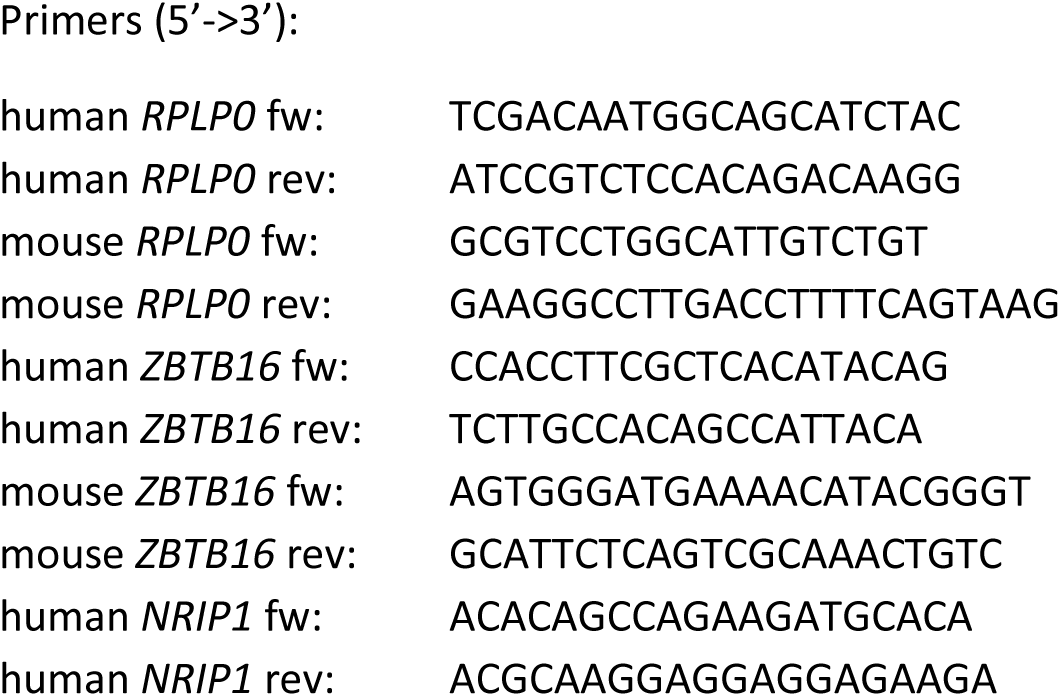

### Functional endothelial cell in vitro assays

In vitro network formation was performed with 1.5×10^5^ HUVECs cultured in 12-well plates (Greiner Bio-One GmbH) that had been coated with 250 µL Matrigel (number 356234; Corning). Network formation was determined after 24 hours by measuring the cumulative tube length in 5 randomly chosen microscopic fields with a computer-assisted microscope using Axiovision 4.5 (Zeiss). Angiogenic sprouting was studied by HUVEC spheroid-based angiogenic sprouting assay as previously described^6^.

Cardiac tissue mimetics (CTMs) containing primary rat cardiomyocytes, fibroblasts and HUVEC, as described previously^4^, were treated with supernatants of siCtrl or siZBTB16-transfected HUVEC for 14d. Beating frequency was observed and CTMs have been stained with Mouse anti-α-smooth muscle actin - Cy3™ (αSMA, 1:200, C6198-2ML, Sigma-Aldrich), 4’,6-diamidino-2-phenylindole (DAPI) and Rabbit anti-COL1A1 (1:100, 72026S, CST).

Cell migration was assessed by Boyden chamber transmigration assays. The assays were conducted in 24-well plates (Corning) using the Boyden chamber (FluoroBlok, 4137, Corning) inserts with 8-µm pores. The bottom chamber was loaded with 800 µL full EBM. 5×10^4^ HUVECs were adjusted in 200 µL and added to the upper chamber. The cells were incubated for 3 hours at 37°C and 5% CO_2_ in a humidified atmosphere, and the filters were fixed in 4% PFA and stained with 4’,6-diamidino-2-phenylindole (DAPI). The number of cells that migrated through the filter was determined by counting the cells in 5 randomly chosen images in the center of the filter at ×10 magnification using the inverted Nikon Eclipse fluorescence microscope.

### Chromatin immuno precipitation (ChIP)

HUVECS expressing HA-tagged *ZBTB16* were fixed with culture medium containing 1% PFA for 10 minutes at room temperature and quenched with 125 mM glycine. The cells were harvested with a scraper and centrifuged at 3000 rpm at 4°C for 5 minutes, and chromatin was isolated with lysis buffer. Isolated chromatin was sheared to fragments of 300-500 bp average length in the Diagenode Bioruptor for 12 cycles with 30-second pulses of ON or OFF at maximum power. 10% of the sheared chromatin was kept as input control, and the rest was added to protein G Dyna beads coated with 5 ug anti-HA antibody (AB9110). Both input and ChIP DNA were treated with RNase, proteinase K, and crosslinks were reversed by incubation overnight at 65 °C followed by phenol-chloroform extraction and ethanol precipitation.

### ChIP sequencing analysis

Trimmomatic version 0.39 was employed to trim reads after a quality drop below a mean of Q15 in a window of 5 nucleotides and keeping only filtered reads longer than 15 nucleotides^7^. Reads were aligned versus Ensembl human genome version hg38 (Ensembl release 109) with STAR 2.7.11a^8^. Alignments were filtered to remove: duplicates with Picard 3.0.0 (Picard: A set of tools (in Java) for working with next generation sequencing data in the BAM format), spliced, multi-mapping, ribosomal, or mitochondrial reads. Peak calling was performed with MUSIC version 20170728 in punctate mode with q-value < 0.2, p-value normalization window length 1500 and enrichment vs. input > 3x^9^. Peaks overlapping ENCODE blacklisted regions (known misassemblies, satellite repeats) were excluded. Remaining peaks were unified to represent a common set of regions for all samples. Counts were produced with featureCounts^10^. The raw count matrix was normalized with DESeq2 version 1.36.0^11^. Peaks were annotated with the promoter (TSS +-5000 nt) of the nearest gene based on Ensembl release 109. Contrasts were created with DESeq2 based on the normalized count peak matrix with all size factors set to one. Peaks were classified as significantly differential at average count > 10 and -1 < log2FC > 1.

### Single nuclei assessment

To isolate nuclei for transcriptomic analysis, hearts were perfused with 1X DPBS in vivo, snap-frozen in liquid nitrogen and stored at -80°C until usage. Eight mice hearts were sequenced (n=4 per group, two females and two males, respectively). For the nuclei isolation, one female and one male heart per group were pooled and processed together. The hearts were minced in pre-filtered homogenization buffer containing 250 mM sucrose (1623637, Sigma-Aldrich), 25 mM KCl (AM9640G, Invitrogen), 5 mM MgCl_2_ (AM9530G, Invitrogen), 10 mM Tris buffer pH 8.0 (AM9855G, Invitrogen), 1 mM DTT (P2325, Thermo-Fisher Scientific), 1X protease inhibitor (11697498001, Roche), 0.6 U/µL Ambion RNase inhibitor (AM2682, Thermo-Fisher Scientific), 0.1% Triton (T8787, Sigma-Aldrich) and 2% BSA (A8022, Sigma-Aldrich) in ultra-pure DNase/RNase-free distilled water (10977035, Thermo-Fisher Scientific). Cells were disrupted with seven strokes of a loose pestle in a glass dounce homogenizer. The homogenized solution was filtered using a pre-wetted 40 µm cell strainer followed by a pre-wetted 20 µm cell strainer. After centrifugation (500 g at 4°C for 8 minutes), the cell pellet was re-suspended in sorting buffer containing 2% BSA, 0.6 U/µL Ambion RNase inhibitor, 1 mM DTT in DPBS. 7AAD (420403, Biolegend) positive nuclei were separated from cell debris using the FACS-Aria III instrument (BD Biosciences; Nozzle size 100 microns) into collection buffer containing 2% BSA, 1 U/µL Ambion RNase inhibitor, 1 mM DTT in DPBS. Sorted nuclei were washed with 0.04% BSA in DPBS.

Library preparation was performed as previously described^12^ and single-nucleus RNA sequencing results were processed with the 10X Genomics Cell Ranger pipeline (7.2.0). The command “*count*” was employed to aligned the raw reads to the mouse reference genome (GRCm38-2019). Following the best practices for single-nucleus RNA sequencing, the option “include-introns” was activated during the analysis. To mitigate ambient RNA contamination, the ‘*remove-background*’ module of CellBender (0.3.0) was applied to the raw data counts^13^. The downstream processing and analysis were conducted with Scanpy (1.9.6)^14^ operated within Python 3.9.18. Our downstream analysis excluded genes expressed in less than 3 cells. We further removed cells with insufficient gene counts (<250 UMI) and with an excess of mitochondrial RNA counts (>10%). Doublet detection was performed using Scrublet (0.2.3) assuming an expected doublet rate of 6%. Additionally, the top 5% of cells with the highest number of UMI counts were discarded. After normalization of the data to total counts and a log-transformation, the most variable genes across the different batches were identified to performed dimensionality reduction and integration through principal component analysis and BBKNN batch correction approach (1.6.0) using 3 neighboring cells within batches. The uniform manifold approximation and projection (UMAP) embeddings were computed to visualize the data. Cell clustering was conducted using the *Leiden* algorithm with a resolution parameter of 0.3. Identified clusters were automatically annotated by using *CellTypist* (1.6.2., Human Heart model V1.0.)^15^. The annotation was further cross-checked with gene markers previously identified for the different cell types of the heart^16–18^. To identify differentially expressed genes between the two conditions, the Wilcoxon rank-sum test and the Benjamini-Hochberg correction method was applied using the “*rank_genes_groups”* function of Scanpy.

### Bulk ATAC sequencing

Extraction Protocol: 50.000 cells were used for ATAC library preparation using Tn5 Transposase from Nextera DNA Sample Preparation Kit (Illumina).

Library Construction Protocol: Cells were pelleted and resuspended in 50µl Lysis/Transposition reaction (12.5µl THS-TD-Buffer, 2.5µl Tn5, 5µl 0.1% Digitonin and 30µl water) and was incubated at 37°C for 30min with occasional snap mixing. Following purification of the DNA fragments was done by MinElute PCR Purification Kit (Qiagen). Amplification of Library together with Indexing was performed as described elsewhere (Transposition of native chromatin for fast and sensitive epigenomic profiling of open chromatin, DNA-binding proteins and nucleosome position^19^. Libraries were mixed in equimolar ratios and sequenced on NextSeq2000 platform (Illumina) using P4 flowcell with 2x 36bp paired-end setup.

ATAC-Seq analysis: Trimmomatic version 0.39 was employed to trim reads after a quality drop below a mean of Q15 in a window of 5 nucleotides and keeping only filtered reads longer than 15 nucleotides (Bolger et al., Trimmomatic: a flexible trimmer for Illumina sequence data). Reads were aligned versus Ensembl human genome version hg38 (Ensembl release 109) with STAR 2.7.11b^8^. Alignments were filtered to remove: duplicates with Picard 3.1.1 (Picard: A set of tools (in Java) for working with next generation sequencing data in the BAM format), spliced, multi-mapping, ribosomal, or mitochondrial reads. Peak calling was performed with Macs version 3.0.0a7 with FDR < 0.0001^20^. Peaks overlapping ENCODE blacklisted regions (known misassemblies, satellite repeats) were excluded. Remaining peaks were unified to represent a common set of regions for all samples. Counts were produced with feature Counts^10^. The raw count matrix was normalized with DESeq2 version 1.36.0 (Love et al., Moderated estimation of fold change and dispersion for RNA-Seq data with DESeq2). Peaks were annotated with the promoter (TSS +-5000 nt) of the nearest gene based on Ensembl release 109. Contrasts were created with DESeq2 based on the normalized count peak matrix with all size factors set to one. Peaks were classified as significantly differential at average count > 10 and -1 < log2FC > 1.

DESeq2 DE: P-value based on Wald test that was corrected for multiple testing using the Benjamini and Hochberg method Kobas GSO: P-value based on hypergeometric test that was corrected for multiple testing using the Benjamini and Hochberg method fGSEA: P-value based on multi-level split Monte-Carlo scheme that was corrected for multiple testing using the Benjamini and Hochberg method Pscan TFBS: P-value based on z-test; no multiple testing correction available by default TOBIAS: P-value of differential binding based on 1) fitting a two-component gaussian-mixture model to the log-transformed background distribution of scores to estimate a threshold between unbound/bound binding sites, and 2) estimating the significance threshold of the right-most normal distribution (by a user-defined p-value); no multiple testing correction available by default

### Statistical analysis

Statistical data are represented as mean and error bars indicate the standard error of mean (SEM). Shapiro Wilk test was carried out the normality distribution analysis. To compare two groups, an unpaired two-sided t-test (Gaussian distributed data) or a Mann Whitney U test (non-Gaussian distributed data) was used. For multi-group comparison, an ordinary one-way ANOVA with a post-hoc Tukey test (Gaussian distributed data) or a Kruskal-Wallis test (non-Gaussian distributed data) was used.

