## Supplementary material for "Endothelial expression of ZBTB16 protects against cardiac aging": Suppl. Figures

### Supplementary online data Fig. 1

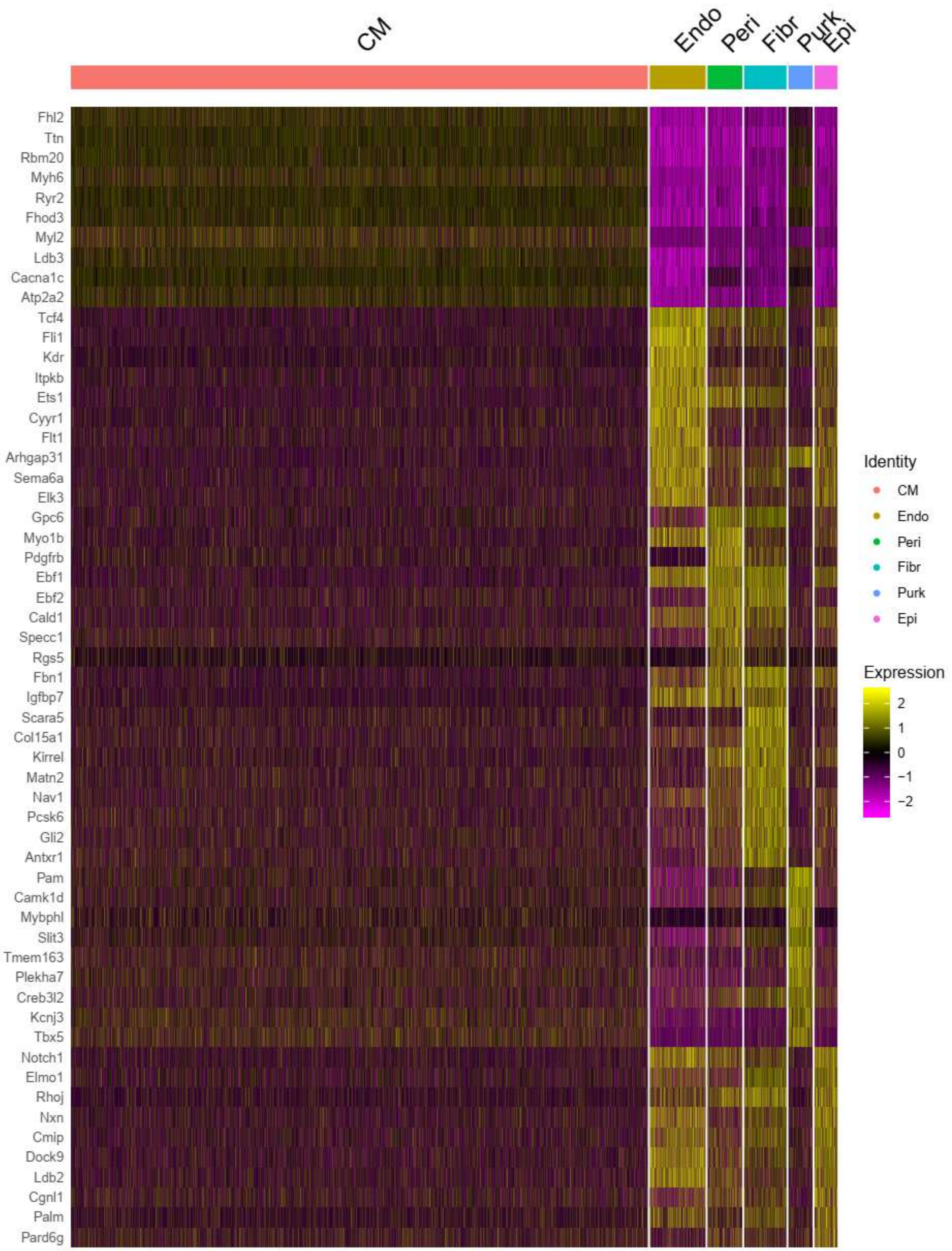


**Supplementary online data Fig. 1: Cell annotation of snATAC-seq data.** Single-nuclei ATAC sequencing (snATAC-seq) was performed on hearts obtained from young (3-month-old) and old (22-month-old) mice (n=3 per group). Shown are the genes used for cell annotation.

### Supplementary online data Fig. 2

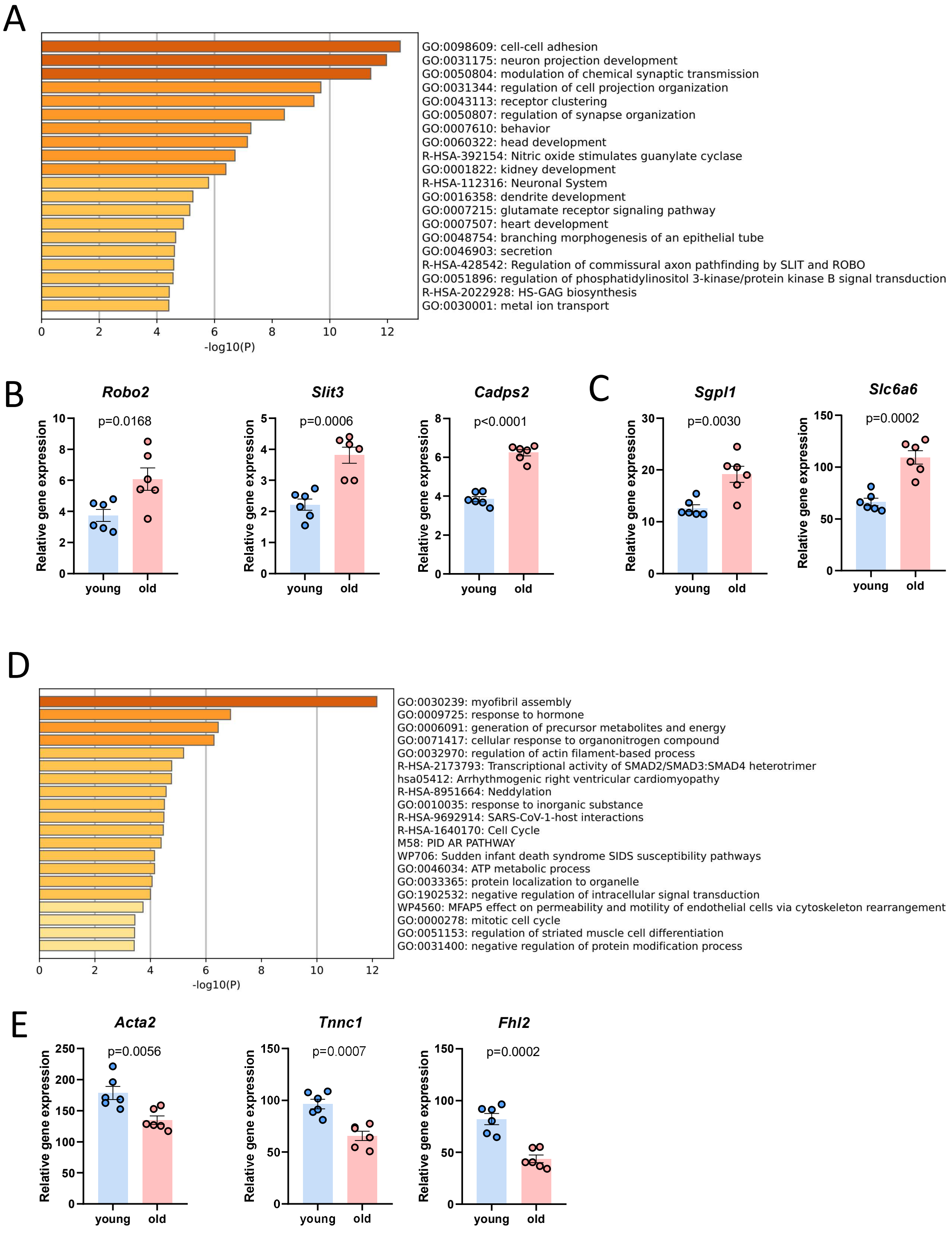


**Supplementary online data Fig. 2: Assessment of differentially accessible genes in aged cardiac endothelial cells.** (A) GO term analyses of genes that were significantly more accessible in aged cardiac endothelia cells (analysis performed with Metascape.org). (B, C) Genes with increased accessibility identified from snATAC-seq were validated in bulk RNA sequencing data from isolated cardiac endothelial cells from young and aged mice (n=6). (D) GO term analyses of genes that were significantly less accessible in aged cardiac endothelia cells (analysis performed with Metascape.org). (E) Genes with decreased accessibility were validated in bulk RNA sequencing data from isolated cardiac endothelial cells from young and aged mice (n=6). Data are shown as mean and error bars indicate the standard error of the mean. P-values were quantified using the unpaired, two-sided t-test.

### Supplementary online data Fig. 3

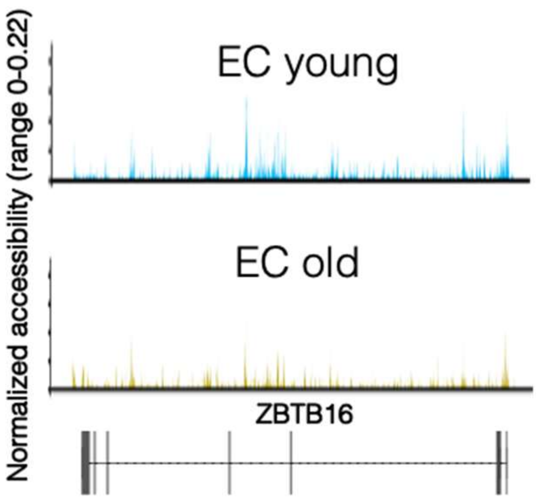


**Supplementary online data Fig. 3: *Zbtb16* accessibility in young and old cardiac endothelial cells.** Representative overview image showing the accessible regions within the *Zbtb16* gene in cardiac endothelial cells of young and old mice.

### Supplementary online data Fig. 4

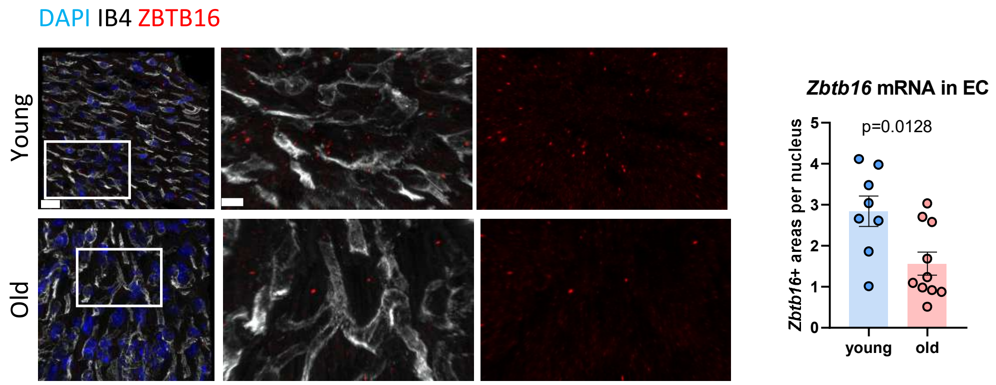


**Supplementary online data Fig. 4: *Zbtb16* mRNA is less abundant in aged cardiac endothelial cells.** Zbtb16 mRNA was visualized in endothelial cells of young (3 months) versus aged (20 months) mouse hearts using RNAscope (red, indicated by yellow arrow heads). IB4 (grey) was used to stain for endothelial cells and DAPI (blue) marks nuclei. Scale bar = 20 µm and 5 µm. Data are shown as mean and error bars indicate the standard error of the mean. P-values were quantified using the unpaired, two-sided t-test.

### Supplementary online data Fig. 5

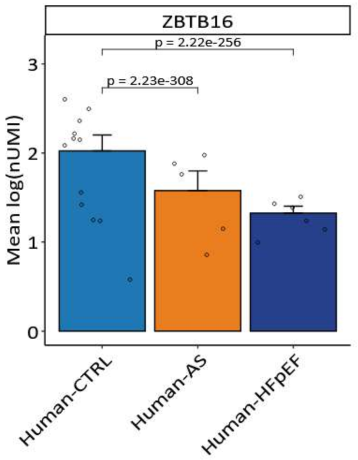


**Supplementary online data Fig. 5: *Zbtb16* expression is decreased in human cardiac cells from aortic stenosis and HFpEF patients.** *Zbtb16* expression was assessed in snRNA sequencing data from heart biopsies derived from cardiac healthy, aortic stenosis (AS) and HFpEF patients. Data are taken from[23,24].

### Supplementary online data Fig. 6

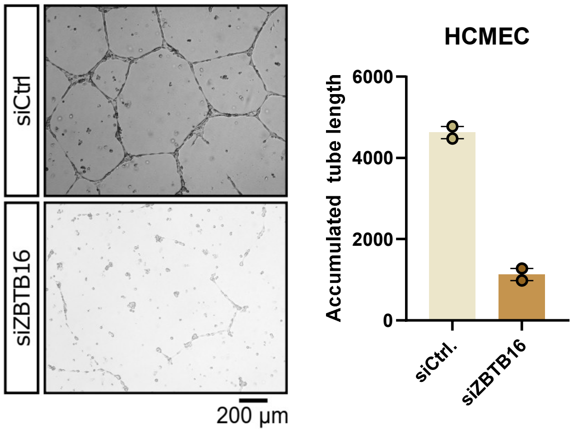


**Supplementary online data Fig. 6: Silencing ZBTB16 reduces network formation in human cardiac microvascular endothelial cells.** To confirm the observations made in HUVEC, the network formation assay was performed as key experiment in primary human cardiac microvascular endothelial cells (HCMEC). Two experimental replicates were performed to prove that ZBTB16 knockdown represses endothelial function in human cardiac specific ECs (n=2).

### Supplementary online data Fig. 7

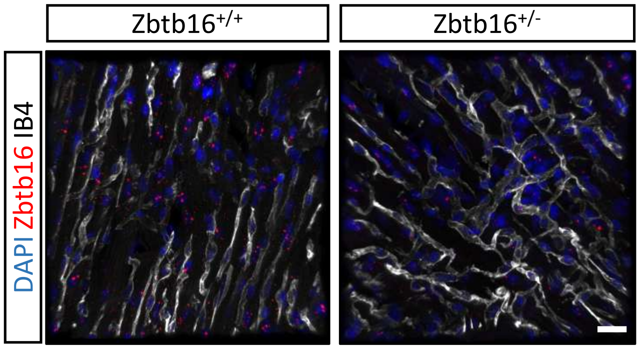


**Supplementary online data Fig. 7: *Zbtb16* deletion in Zbtb16+/- mice.** Representative images of RNA-scope against Zbtb16 mRNA (red), DAPI (blue) and IB4 (grey). Scale bar = 20µm.

### Supplementary online data Fig. 8

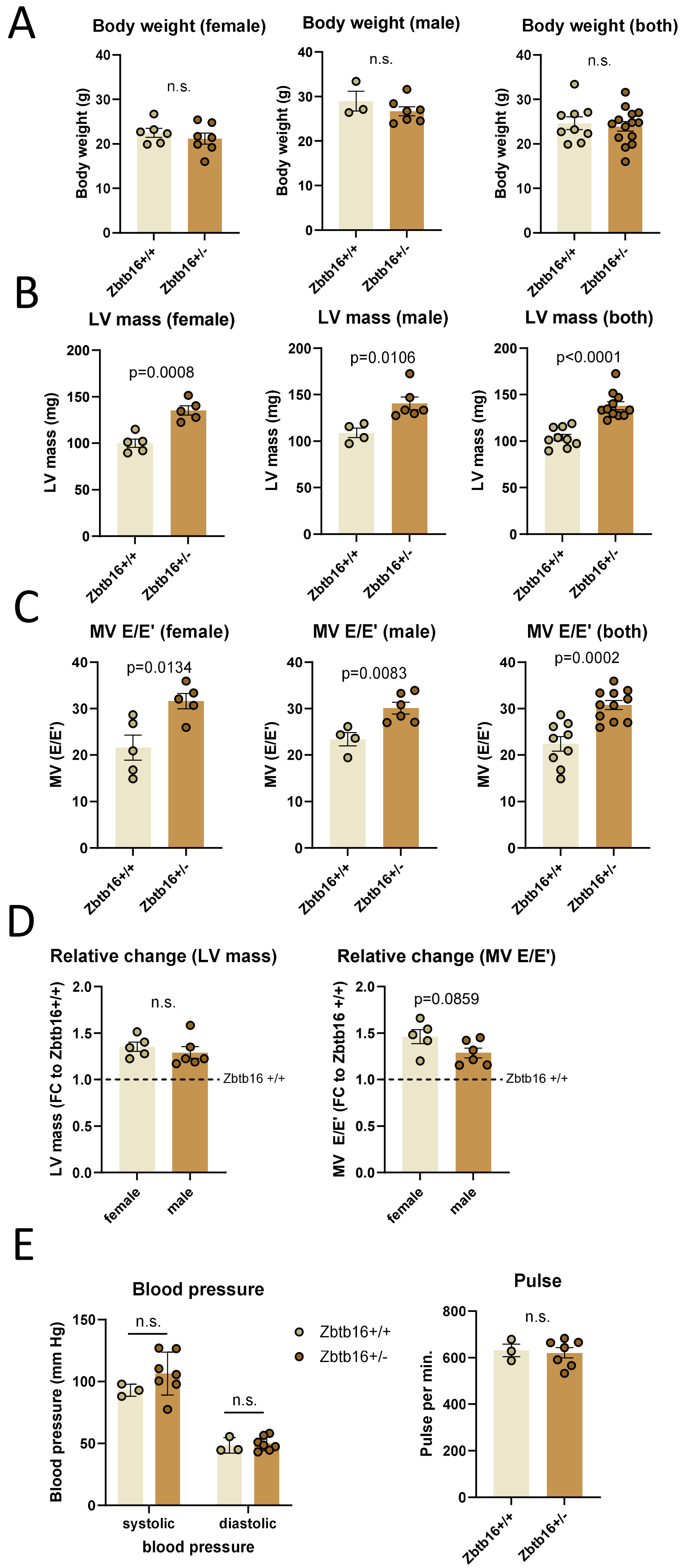

**Supplementary online data Fig. 8: *Zbtb16* deletion in Zbtb16+/- mice.** (A-D) Body weight and echocardiography data assessing the left ventricular (LV) mass (B, D) and diastolic function (MV E/E', C, D) in female (n=5 vs. n=5) and male (n=4 vs. n=6) Zbtb16+/+ versus Zbtb16+/- mice. (E) Tail-cuff blood pressure measurements in Zbtb16+/+ (n=3) versus Zbtb16+/- (n=7) mice. Shown are both systolic and diastolic blood pressure, as well as pulse per minute of these animals. Data are Gaussian distributed and statistical power was assessed using the unpaired, two-sided t-test.

### Supplementary online data Fig. 9

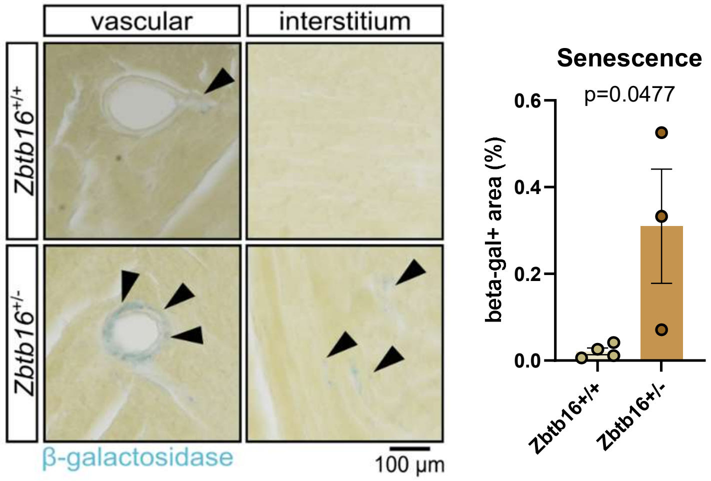


**Supplementary online data Fig. 9: *Zbtb16* deletion induced cardiac senescence.** Hearts of 3- to 4-month-old *Zbtb16*+/+ versus *Zbtb16*+/- (n=4 vs. n=3) were probed for cellular senescence (β-galactosidase, blue). Data are represented as mean and error bars indicate the standard error of the mean. Data are Gaussian distributed and p-value was calculated using the unpaired, two-sided t-test.

### Supplementary online data Fig. 10

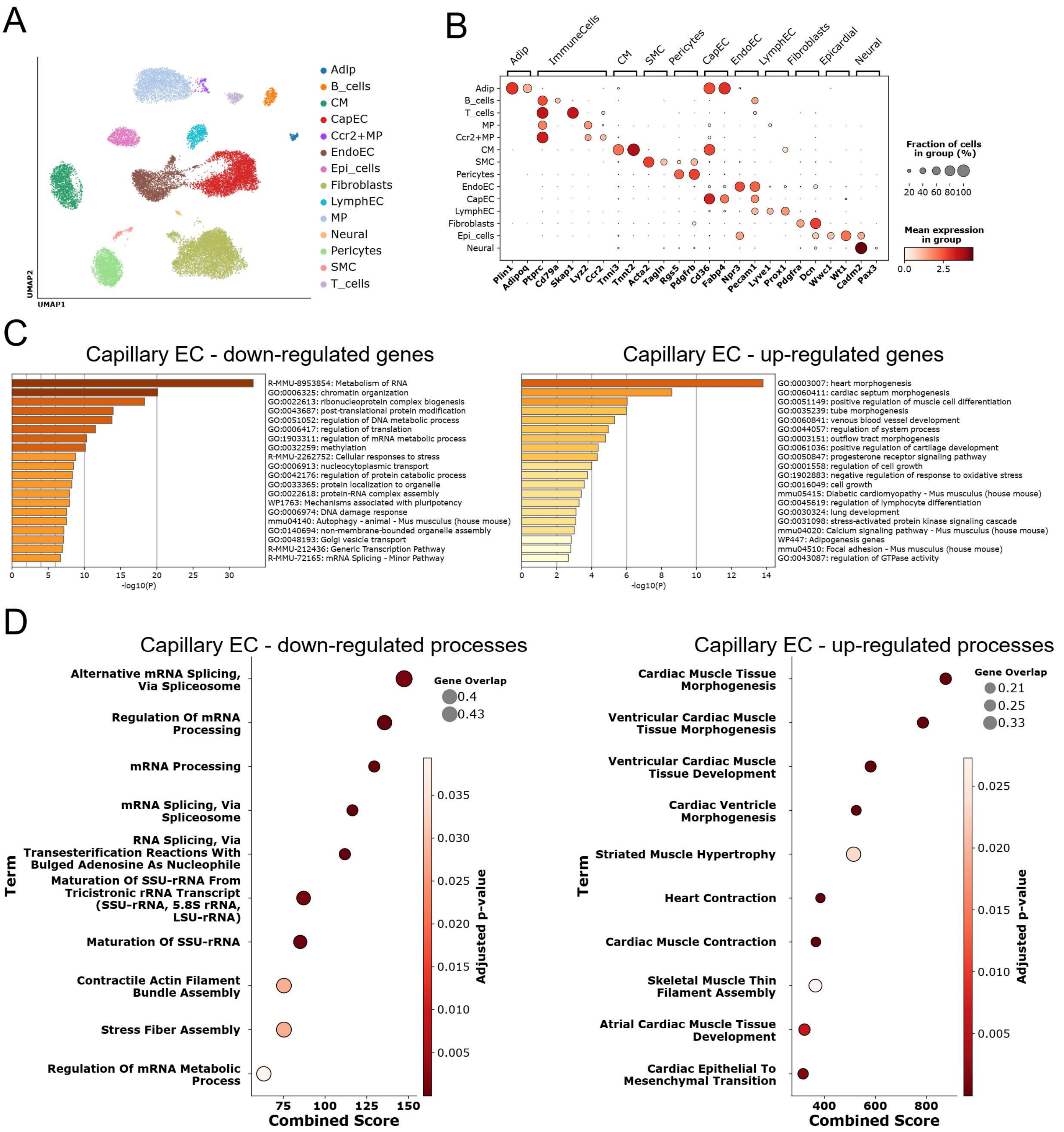


**Supplementary online data Fig. 10: Single-nuclei RNA sequencing reveals transcriptomic changes in cardiac endothelial cells upon *Zbtb16* deletion.** Hearts of 3- to 4-month-old *Zbtb16*+/+ versus *Zbtb16*+/- (n=2 each) were used for snRNA-Seq. (A) UMAP plot displaying annotated cell types. (B) Cell annotation to identify specific cell clusters. (C) GO terms associated with significantly regulated genes of the capillary EC cluster in *Zbtb16*+/- mice as analyzed by Metascape. (D) Biological processes associated with significantly regulated genes of the capillary EC cluster as analyzed by EnrichR.

### Supplementary online data Fig. 11

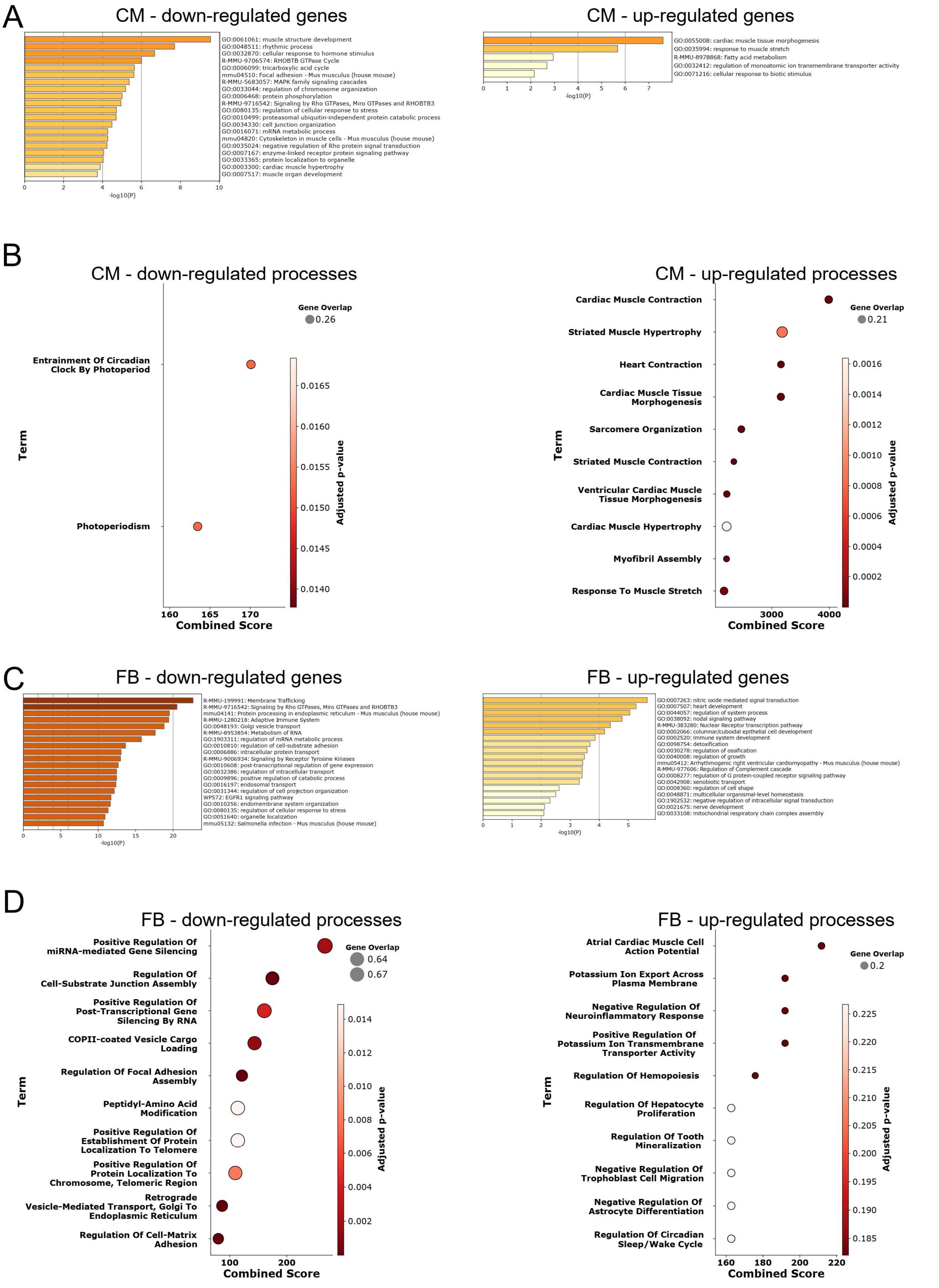

**Supplementary online data Fig. 11: Single-nuclei RNA sequencing reveals transcriptomic changes in cardiac CM and FB upon *Zbtb16* deletion.** Hearts of 3- to 4-month-old *Zbtb16*+/+ versus *Zbtb16*+/- (n=2 each) were used for snRNA-Seq. (A) GO terms associated with significantly regulated genes of the cardiomyocyte (CM) cluster in *Zbtb16*+/- mice as analyzed by Metascape. (B) Biological processes associated with significantly regulated genes of the cardiomyocyte (CM) cluster as analyzed by EnrichR. (C) GO terms associated with significantly regulated genes of the fibroblast (FB) cluster in *Zbtb16*+/- mice as analyzed by Metascape. (D) Biological processes associated with significantly regulated genes of the fibroblast (FB) cluster as analyzed by EnrichR.

### Supplementary online data Fig. 12

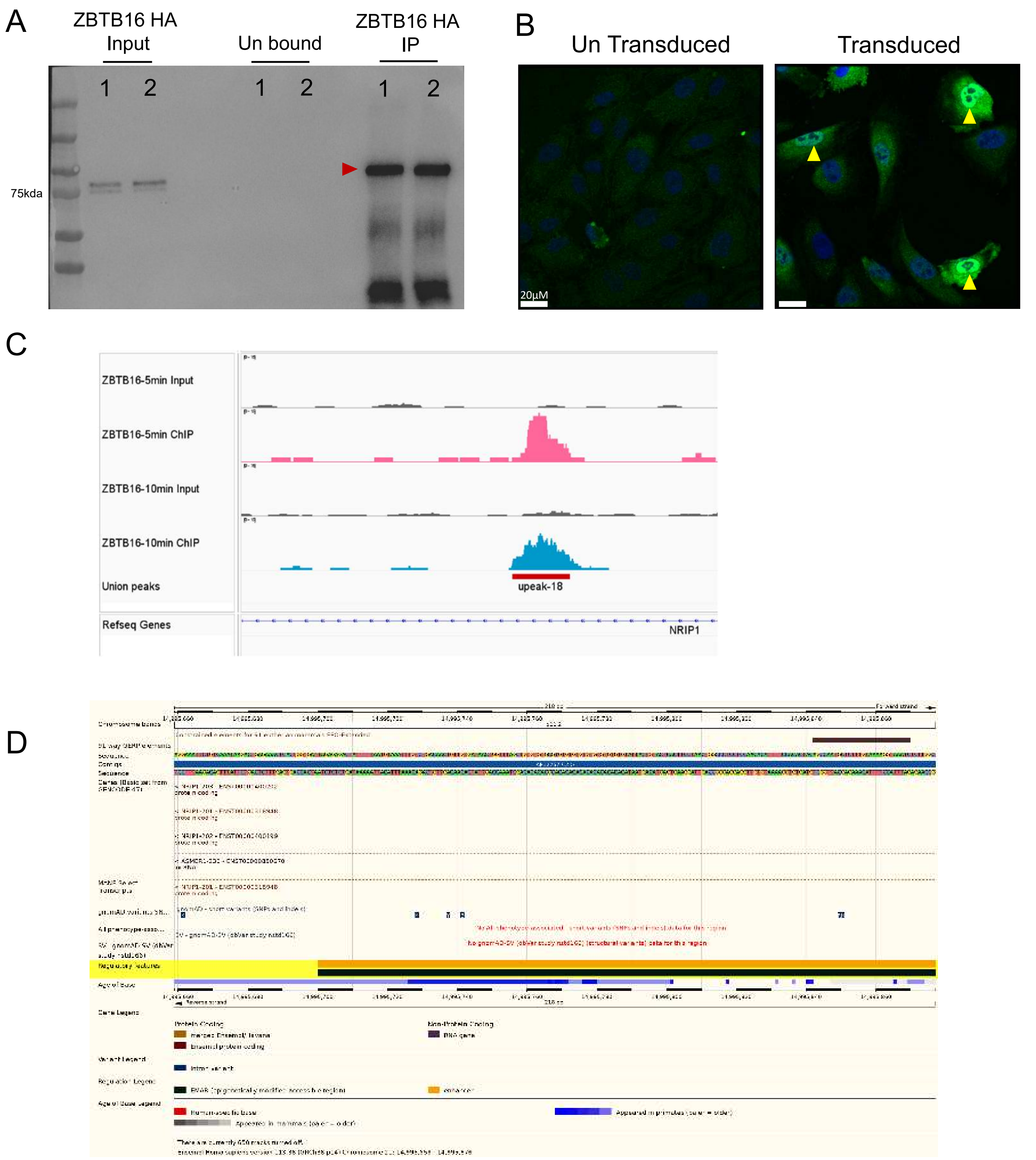


**Supplementary online data Fig. 12: Identification of NRIP1 as ZBTB16 target.** (A) Immunoprecipitation of HA-tagged ZBTB16 with HA antibody, numbers indicate independent lentiviral transduction and red arrows indicate the enrichment of HA-tagged ZBTB16. (B) Immunofluorescence staining for HA tag on HUVECS transduced with HA-tagged ZBTB16. (C) Representative images of the peaks on NRIP1 on ZBTB16 ChIP after 5 or 10 min crosslinking. (D) localization of the peak in the NRIP gene.

### Supplementary online data Fig. 13

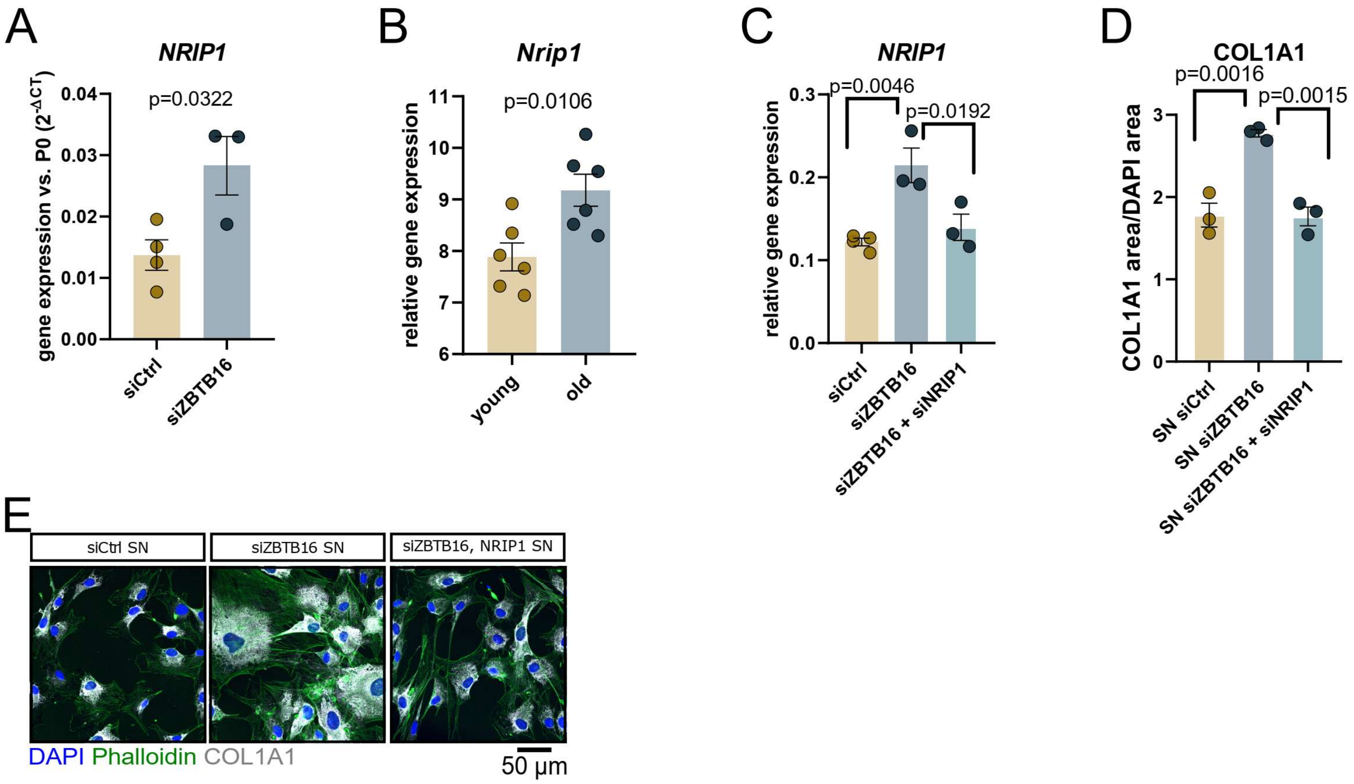


**Supplementary online data Fig. 13: ZBTB16 repression of NRIP1 contributes to paracrine activation of fibroblasts** (A) *Nrip1* gene expression in cardiac endothelial cells isolated from young and old mouse hearts (n=6). Data taken from [8]. (B) *NRIP1* gene expression in HUVEC transfected with non-targeting siRNAs (siCtrl, n=4) versus siRNAs against *ZBTB16* (siZBTB16, n=3). (C) *NRIP1* gene expression in HUVEC transfected with non-targeting siRNAs (siCtrl, n=4) versus siRNAs against *ZBTB16* (siZBTB16, n=3) versus siRNAs against *ZBTB16* and *NRIP1* (siZBTB16+siNRIP1, n=3). (D-E) HUVEC were transfected with either non-targeting siRNAs (siCtrl), siRNAs against ZBTB16 (siZBTB16) or with siRNAs against ZBTB16 and NRIP1 (siZBTB16+siNRIP1). Supernatants were collected and transferred to human cardiac fibroblasts (n=3). After 72h, fibroblasts were stained for COL1A1 (grey), DAPI (blue) and Phalloidin (green). Quantification of COL1A1 is shown in D, representative images in (E). Data are shown as mean and error bar indicate the standard error of the mean. P-value were calculated by two-tailed Student’s t-test (A, B) or by using an ordinary one-way ANOVA with a post-hoc Tukey’s test (C, D).

### Supplementary online data Fig. 14

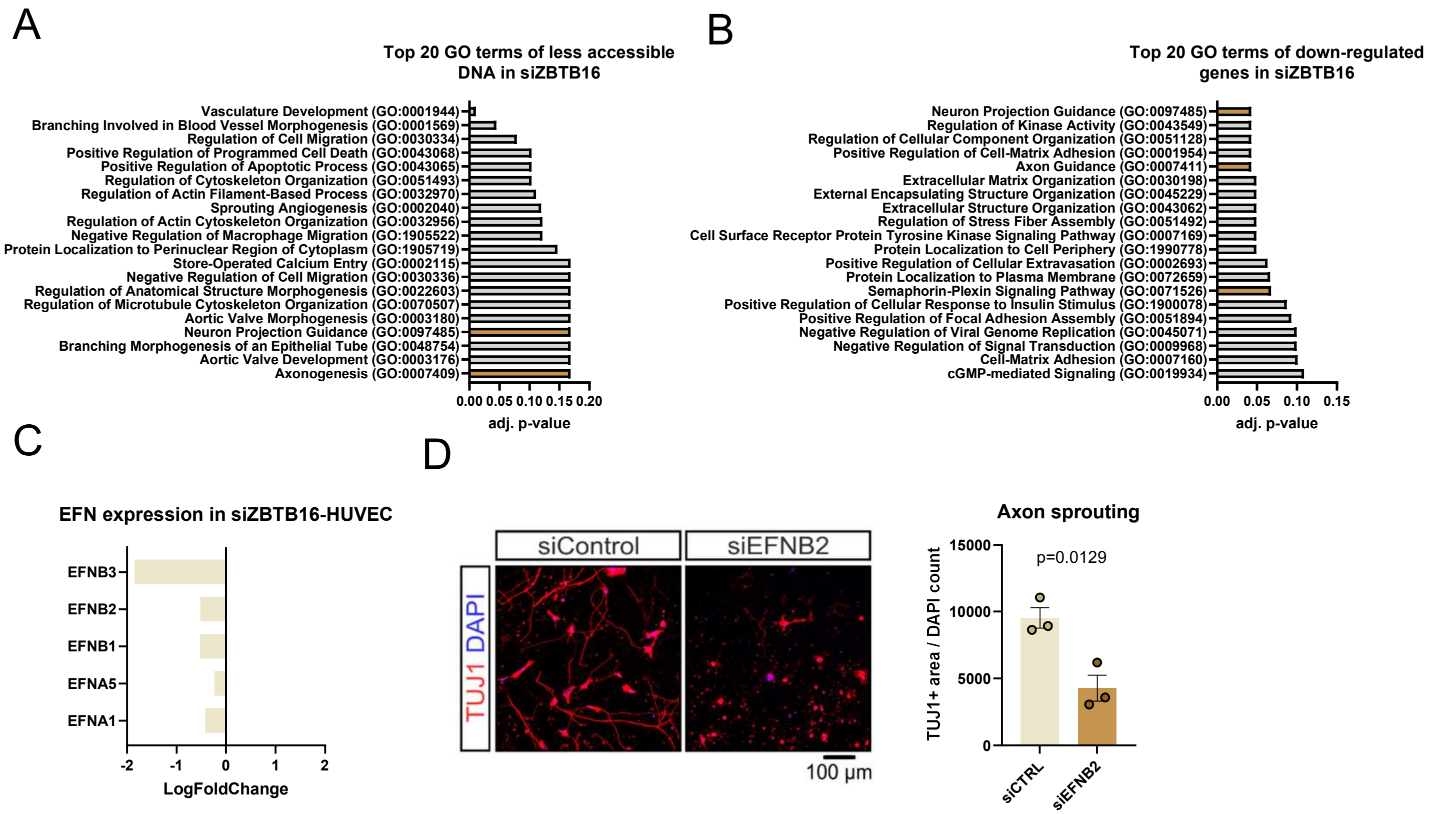


**Supplementary online data Fig. 14: ZBTB16 regulates endothelial-nervous cross-talk via ephrins.** (A) Bulk ATAC sequencing of HUVEC after siZBTB16 or control transfection (n=3). GO terms of less accessible DNA regions are shown. (B) GO term analysis of significantly down-regulated genes in siZBTB16 treated HUVEC. GO terms were assessed with Enrichr. (C) EFN family member expression (Log2 FC) in HUVEC after siZBTB16 treatment (n=5). (D) HUVEC were transfected with either non-targeting (siCtrl) or EFNB2 targeting siRNAs (siEFNB2). Supernatants were collected and transferred to mouse cortical neurons. Sprouting was assessed by TUJ1 and DAPI staining. (n=3). Data are Gaussian distributed and p-values were calculated using two-sided, unpaired t-test.

### Supplementary online data Fig. 15

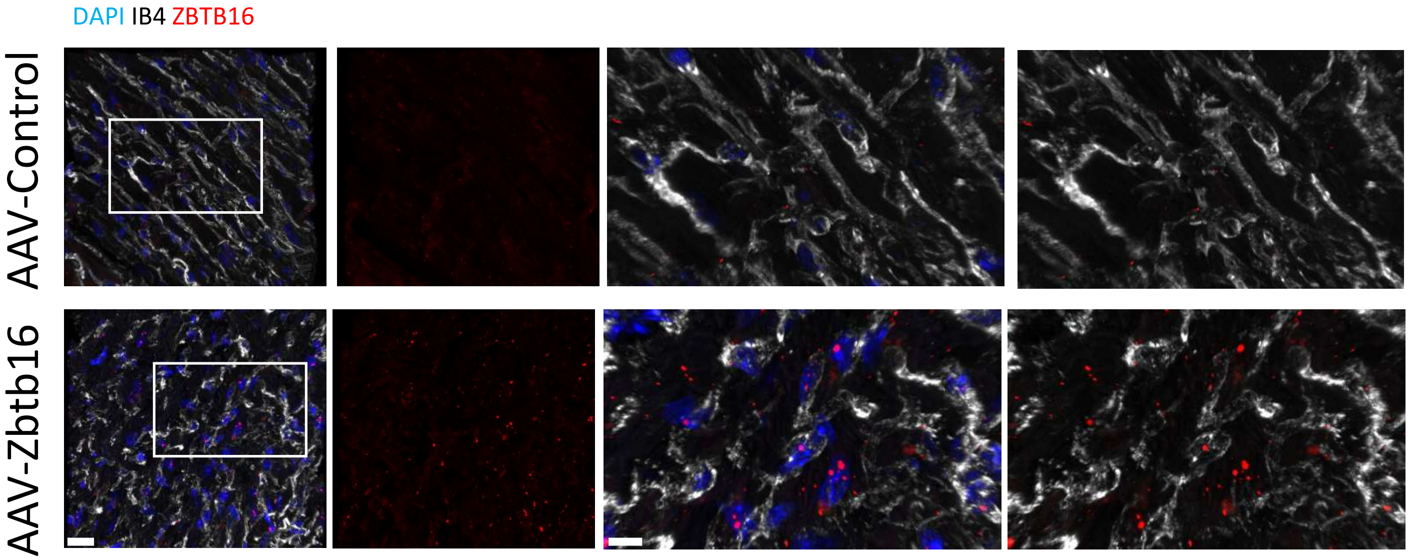


**Supplementary online data Fig. 15: AAV9-mediated *Zbtb16* overexpression in cardiac endothelial cells.** *Zbtb16* was overexpressed in endothelial cells of 18-month-ol C57Bl/6J mice. Eight weeks after AAV9 treatment, heart sections were stained for *Zbtb16* mRNA using RNA-scope (red). Isolectin B4 (grey) stains endothelial cells and DAPI marks nuclei (blue).

### Supplementary online data Fig. 16

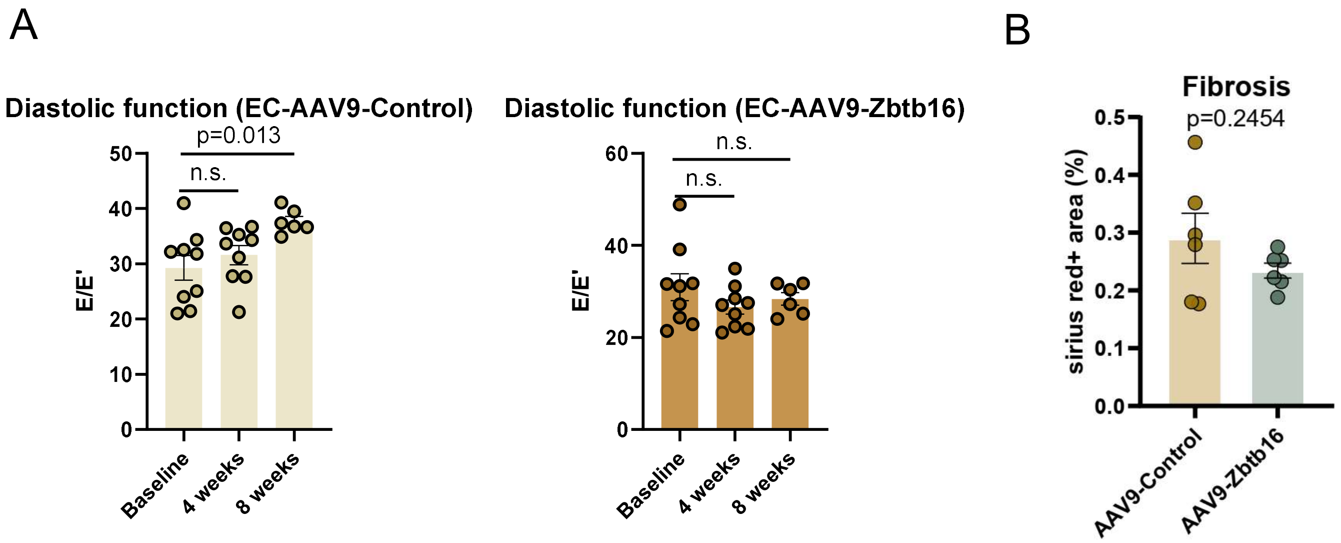


**Supplementary online data Fig. 16: AAV9-mediated *Zbtb16* overexpression in cardiac endothelial cells attenuates interstitial fibrosis.** (**A-B**) Overexpression of Zbtb16 in endothelial cells by targeted AAV9 vectors improves cardiac function in 18 month old mice. (A) Diastolic function (E/E') as assessed by echocardiography at baseline and after 4 and 8 weeks after AAV9 treatment (n=9). (B) Sirius red staininig of cardiac sections 8 weeks after AAV9 treatment (n=6). Data are shown as mean and error bar indicate the standard error of the mean. P-value was calculated by two-tailed Student's t-test (B) or by One-way ANOVA with a post-hoc Dunnett's test (A).
